# Modification of RNAs in natural rG4 (MoRNiNG), A Database of RNA Modification Sites Associated with the Dynamics of RNA Secondary Structures

**DOI:** 10.1101/2025.09.11.675469

**Authors:** Yicen Zhou, Shanxin Lyu, Shiau Wei Liew, Xi Mou, Ian Hoffecker, Jian Yan, Yu Li, Chun Kit Kwok, Jilin Zhang

## Abstract

RNA structures are essential building blocks of functional RNA molecules. Profiling secondary structures *in vivo* and in real-time remains challenging because RNAs exhibit dynamic structures and complex conformations. Besides the stem-loop canonical secondary structure, the non-canonical structure RNA G-quadruplex (rG4) is of great interest for its potential as a drug target. Early studies have demonstrated that RNAs can form distinct secondary structures. However, it is not well understood how distinct RNA structures, formed from the same RNA sequences, function within the transcriptome. The factors driving and regulating structure transitions remain poorly investigated. Inspired by a segment in the *HOXB9* mRNA capable of forming multiple structures, we found that many RNA segments across the transcriptome exhibit multi-faceted structure-forming potential. For the *HOXB9* case, we demonstrated that RNA modification influences RNA structure and binding with RNA binding proteins. Therefore, we collected RNA modification sites naturally occurring in the putative G-quadruplex-forming sequences (PQSs) of transcripts and developed MoRNiNG, a freely accessible database at https://www.cityu.edu.hk/bms/morning. The database is designed and organized by reliability tiers determined by the resolution of RNA modification sites and capable of including various large datasets. We experimentally validated the influence of m6A, m5C, and A-to-I editing on rG4-forming sequences and provided evidence to support the modification switch concept. The diversity and transition of secondary structures from the same RNA segment offers valuable insights into the regulation of RNA structure dynamics.

## Introduction

RNA molecules are instrumental in many biological processes [1]. Dissecting their functional roles is crucial in constructing a comprehensive regulatory map. One of the significant challenges hindering this process is the elusive information that determines the transition between linear sequences and structural RNAs, as well as between different RNA structures.

RNAs form a variety of structures, and a growing volume of RNA profiling data has demonstrated that many bases participate in canonical base pairing [2, 3]. Beyond canonical secondary RNA conformations, such as stem-loop (SL) structure, internal loops and multi-branch junctions, RNA can form the non-canonical structure RNA G-quadruplex (rG4) by looping back to stack more than two planar G-tetrads, each consisting of four guanines connected via Hoogsteen hydrogen bonds and stabilized by a monovalent cation [4, 5]. Since its discovery, the rG4 structure has gained recognition as a potential functional element in both transcriptional and post-transcriptional regulation. Numerous assays have been devised and refined to profile rG4 *in vitro* and *in vivo* [4, 5], offering unprecedented transcriptome-wide insight into rG4 distribution and functional. However, the rG4 landscape rG4 dynamics require substantial exploration to gain a comprehensive understanding.

The flexible nature of single-stranded RNAs allows them to dynamically change conformation to interact with RNA binding proteins with structural specificity [6, 7]. A typical *in vivo* example is LIN28A, which binds to the pre-let-7 (preE) terminal SL structure through its cold shock domain to inhibit the function of miRNA *let-7* [8]. This protein can also unwind rG4 in mRNA to promote ribosomal scanning and enhance translation [9]. The polycomb repressive complex 2 (PRC2) interacts with rG4 in *Xist,* which is crucial to silence genes on chromosome X in a stepwise manner [10]. These cases emphasize the importance of structural RNA in regulating gene expression. In addition to previously reported synthetic ligands, evidence suggests that a single RNA sequence can adopt distinct structures, indicating that the biological function of RNA *in vivo* is related to structural specificity [11, 12]. Moreover, the N6-methyladenosine (m6A) modification at the binding site of LIN28A on *HOXB9* mRNA lowers LIN28A’s binding affinity, suggesting a potential alteration of RNA secondary structure. As RNA structures across cellular states remain incomplete and the factors driving conformation are still poorly understood, further investigation is needed to elucidate transitions between secondary structures.

RNA modifications, such as m6A, are chemical changes that can alter the nature of local RNA sequences and corresponding RNA structure [13–16]. Indeed, these modifications play vital roles in regulating the formation and stability of rG4 [17, 18]. In the case of miRNA *let-7*, the modification 7-methylguanine (m7G) changes the SL to rG4 equilibrium to allow for DROSHA binding to promote the miRNA maturation [19]. Notably, the key component of methyltransferase, METTL14, prefers binding to rG4 [18], underscoring the role of rG4 in regulating m6A-dependent biological processes. Yet, the impact of RNA modification on the RNA secondary structures remains poorly characterized and requires further investigation. Despite the need to increase the spectrum and resolution of the RNA modification, gaining a large volume of rG4 profiles still requires substantial work. The emerging consequences of RNA modifications in reshaping RNA structures should also be considered to gain better insights into RNA function.

Therefore, to bridge the knowledge gap between RNA epigenetics and structure dynamics, we present a curated compendium of rG4-forming sequences with various RNA modification sites observed in cells that are partially supported by experimental evidence. Although rG4 diversity is not clearly defined at this stage, we have provided evidence to support the observation that RNA modifications can modulate the rG4 structure. The database design is structured to accommodate future extension and ensure compatibility with various data types, laying the foundation for a more comprehensive understanding of RNA structure dynamics.

## Materials and methods

### Oligonucleotides and G4 binding small molecules

The original human *HOXB9* sequence and its derivatives were synthesized with and without FAM tag on the 5’ end from Sangon and Integrated DNA Technologies (IDT), respectively. They were dissolved to a concentration of 100 μM (according to supplier instructions) with ultra-pure nuclease-free distilled water (10977015, Thermo, Waltham, MA). All the dissolved oligos were stored at -20℃ before the experiment. Thioflavin T (ThT, HY-D0218) and N-methyl Mesoporphyrin IX (NMM, HY-133821) were ordered from MedChemExpress (Shanghai, China). SYBR Gold was ordered from Invitrogen (S11494, Waltham, MA).

### ThT and SYBR Gold native gel staining of rG4

Reaction mixtures containing 10 mM Tris-HCl pH 7.5, 150 mM KCl, and 5 μM oligonucleotide were prepared in a 20 μL reaction. The mixtures were heated at 95 ℃ for 5 minutes for denaturation and cooled to room temperature for 15 minutes. 3 µL of 40% sucrose (final ∼5%) was added to the mixtures for gel loading. The mixtures were loaded into a 10% native polyacrylamide gel (Acr-Bis 19:1) with 5-10 µL of volume per well. The gel was run at 300 V at 4 ℃ for 20 minutes. For ThT staining gel, the gel was stained with 0.32 µg/mL (1 μM final) ThT for 5 minutes by adding 5 µL of 10 mM ThT and 50 mL of milli-Q water. For SYBR Gold staining, the gel was stained with 1× SYBR Gold for 5 minutes by adding 5 µl of 10000 × SYBR Gold and 50 ml of milli-Q water. The gels were scanned with SYBR Gold mode using Biorad ChemDoc imaging system (Hercules, CA).

### Ligand-enhanced fluorescence assay

Sample solutions containing 0.5 μM RNA were prepared in 150 mM LiCl/KCl and 0.5 or 1μM ligand (NMM/ThT). Fluorescence spectroscopy was performed using a Molecular Devices SpectraMax ID5 microplate reader, and a 96-well plate was used with a sample volume of 150 μL. Before the measurement, the samples (ligand not added) were denatured at 95 ℃ for 3 minutes and allowed to cool at room temperature for 15 minutes. The samples were excited at 395 nm for NMM and 425 nm for ThT. The emission spectra were acquired from 550 to 750 nm and 465 (or 485) to 650 nm for NMM and ThT, respectively. Data were collected every 1 nm at 25 ℃.

### Circular dichroism (CD) spectroscopy

CD assay was performed with a Jasco CD J-150 spectrometer and a 1 cm path length quartz cuvette. The reaction samples containing 5 μM oligos were prepared in 10 mM LiCac buffer pH 7.0 and 150 mM KCl/LiCl to form a total volume of 2 mL. The mixtures were then vortexed and heated at 95 ℃ for 5 minutes and cooled down to room temperature for 15 minutes for renaturation. The samples were scanned at a 2 nm interval from 220 to 310 nm, and the data were blanked and normalized to mean residue ellipticity before being smoothed over 8 nm.

### UV melting assay

UV melting assay was performed with a Cary 100 UV–vis spectrophotometer and a 1 cm path length quartz cuvette. The reaction samples containing 5 μM oligos were separately prepared in 10 mM LiCac buffer pH 7.0 and 150 mM KCl to form a total volume of 2 ml. The mixtures were vortexed and heated at 95 °C for 5 minutes and allowed to cool for 15 minutes at room temperature for renaturation. In cuvettes sealed with Teflon tape, the samples were monitored at 295 nm from 20 °C to 95 °C with data collected every 0.5 °C. The data were blanked and smoothed over every 10 °C.

### LIN28A protein purification

The *LIN28A* gene and His-tag were inserted into the pET-28ɑ vector and then cloned in *E. coli* BL21 (DE3). Bacteria were cultured in Luria Bertani (LB) broth medium with 100 μg/mL kanamycin overnight at 37 °C on a shaker. LB included 0.5% (w/v) yeast extract, 1.0% (w/v) Tryptone and 0.5% (w/v) NaCl in distilled water (pH 7.2). The culture was inoculated (1:100) into a fresh LB broth until OD600 = 0.5–0.6 was reached. To induce protein synthesis, 0.25 mM IPTG was added to the culture, which was then shaken overnight at 37 °C. Cells were collected by centrifugation at 4000 rpm for 30 minutes. Then, cells were lysed and disrupted by 3 cycles of freezing (-80 °C), thawing (37 °C), and sonication. Afterward, the mixture was centrifuged at 4000 × g for 20 min, and the clear supernatant was passed through a 0.45 μm filter. Subsequently, protein purification was performed using BioRad Ni-charged resin (1560125, Hercules, CA) for His-tagged recombinant proteins (in the denaturing condition) according to the manufacturers’ recommendation. Protein purity was evaluated by 10% SDS-PAGE. Finally, the concentration of proteins was determined by Nanodrop assay.

### Electrophoretic mobility shift assay (EMSA)

RNAs were heated at 95 ℃ for 3 minutes for denaturation and cooled to room temperature for 15 minutes. Reaction mixtures containing 10 nM 5’ FAM-labeled RNA, 20 mM HEPES pH 7.5, 100 mM KCl (stable rG4) or 100 mM LiCl (unstable rG4), 0.12 mM EDTA, 25 mM Tris– HCl (pH 7.5), 10% glycerol, and increasing concentrations of LIN28A were prepared in 10 μL and incubated at 37 ℃ for 1 h. The RNA-protein mixture was resolved by 10%, 37.5:1 (acrylamide: bis-acrylamide) non-denaturing polyacrylamide gel in 0.5 × Tris/Borate/EDTA (TBE) at 4 ℃, under the setting of 100 V for 30 minutes or 150 V for 40 minutes.

#### SL structure prediction and icSHAPE reactivity score distribution of the *HOXB9* ligand

The icSHAPE reactivity scores of *HOXB9* sequence were downloaded from RASP database, using the experiments conducted in HEK293T and HEK293 cell lines [20]. The data were grouped into *in vivo* and *in vitro* categories according to the experimental methods. The icSHAPE scores were subjected to RNAfold to predict the secondary structure [21]. The heatmap was visualized using the ComplexHeatmap [22]. The score of guanines in G-tetrad and other bases from the above six samples was examined.

### icSHAPE reactivity score distribution of PQSs supported by rG4-seq and non-PQSs

For human PQSs supported by rG4-seq, only those containing at least 1nt in any loop and no bulges/mismatches in G-tetrads were considered. The corresponding imputed icSHAPE scores of H9, HEK293, HeLa, K562, and HepG2 cell lines were extracted from the RASP database [20]. PQSs with null reactivity scores at any position were further excluded. Scores of bases within G-tetrads were compared with those of bases in loops. Additionally, an equal number of non-PQS RNA fragments with the same length as sampled PQSs were randomly selected from HEK293 and HeLa cells to compare the score of guanines. The score difference between groups was examined by the Mann-Whitney test. Statistical significance was defined as follows: \**P* < 0.05, \*\**P* < 0.01, \*\*\**P* < 0.001, and \*\*\*\**P* < 0.0001.

### Annotation of rG4 and PQS

Three types of rG4 datasets were included to gain transcriptome-wide rG4 features or potential rG4 loci. This includes the *in vivo* experimental data, rG4-seq and ultra-low-input rG4-seq [23–26], *in vitro* CD spectrum, and rG4 prediction results generated by several tools: pqsfinder [27], QGRS [28], G4Hunter (G4H) [29], G4RNA screener [30], RNAfold [21], G4 Neural Network (G4NN) score > 0.5 and Quadparser-based (G_≥2_N_1–12_)_3_G_≥2_ sequence motif (G_2_L_12_ motif). As the detecting capacity and scope of *in vivo* approaches are usually restricted, we applied a hierarchical predicting scheme to gain consensus rG4 loci to strengthen the reliability and control false-positive loci.

PQSs were primarily predicted within the longest transcript through pqsfinder (minimum score = 1) and QGRS (default parameters). The candidate PQSs were processed by a customized Python script to extract the consensus PQSs, which were further fed into G4Hunter, G4RNA screener, and RNAfold to increase the reliability. In addition to the core PQSs predicted by the two software programs, regions were labeled as ‘G4Hunter score ≥ 1.2’ for PQS or rG4 where G4Hunter results met the parameter settings ‘-w 25 -s 1.2’. These regions were further annotated with the supporting source ‘G4RNA Screener’, but only if the Consecutive G over consecutive C ratio (cGcC), G4H, and G4NN scores were greater than or equal to 4.5, 0.9, and 0.5, respectively. RNAfold was additionally employed to examine whether PQSs could form rG4. One PQS was considered as experimentally supported if it was completely covered by rG4-seq peaks from HeLa cells or if the terminal G of the PQS was located either at the RTS site or 1nt upstream of rG4-seq2.0 of HEK293T cells [24] or rG4-seq of HeLa cells [25]. PQSs on mouse transcripts were additionally examined by intersecting with rG4 peaks identified by ultra-low-input rG4-seq experiments [26]. Putative GI-quadruplex-forming sequences (PIQSs), a type of G-quartet induced by substituting guanine with inosine through RNA editing in the human transcriptome, were also included [31]. As the predicted PIQSs by pqsfinder (score ≥ 10) were not all from the longest transcripts, a separate “modType” option was introduced on the search page.

The distribution of all PQSs on the mRNAs and lncRNAs was visualized using Guitar R package [32], with the annotation of 5’ UTR, CDS, and 3’ UTR regions. The number of PQS in the transcript is obtained by the following formula: normalized count = (PQS count × 1000)/transcript length. The kernel density estimation was performed using a Gaussian kernel.

### PQSs in eCLIP peaks

The PQSs within the longest mRNA and lncRNA transcripts of human transcriptome were intersected with eCLIP data (GRCh38) of K562 and HepG2 cell lines from the ENCODE database [33]. First, the peaks of distinct replicates were merged for each RNA binding protein (RBP) with bedtools [34]. The merged peaks were then intersected with PQS coordinates to obtain the peaks covering at least 50% of nucleotides in the overlapping PQSs. After removing duplicated PQSs, the proportion of PQSs supported by eCLIP peaks in the overall PQSs was calculated.

### Gene ontology enrichment analysis

To explore the functional roles of potential rG4 binding RBPs, transposable elements (TEs) - free eCLIP peaks overlapping with PQSs were defined as rG4 binding sites for each RBP in HepG2 and K562 cell lines. The number of rG4-RBP peaks was then divided by the total count of non-TE peaks to get the ratio of rG4-peaks for each RBP. The gene ontology (GO) enrichment analysis was performed on RBPs with a rG4-peak ratio higher than 15% using clusterProfiler [35]. In total, 41 and 57 rG4 binding RBPs were defined in HepG2 and K562 cells, respectively.

### PQSs supported by rG4-seq

PQSs supported by rG4-seq were identified using the following criteria: PQSs completely covered by rG4-seq peaks or with the last G falling into the rG4-seq and rG4-seq2.0 RTS sites or 1nt upstream. In the human transcriptome, PQSs supported by rG4-seq were counted by merging evidence from all available experimental sources, considering the low count of cell-specific peaks. For the PQS presenting RNA modification, only consistent sources for both RNA modification and rG4-seq (HeLa) and rG4-seq2.0 (HEK293T) were considered.

### Prediction of SL structure embedded in PQSs

The PQS in the longest transcripts, including mRNA, lncRNA from the Ensembl database, miRNA shared by Ensembl and miRBase databases, and all remaining RNA types (Others), were subjected to RNAfold to predict the canonical SL structure [21].

### Comparison of PQSs with public G4 databases

CD-HIT-EST [36] was used to cluster the 178,974 unique human PQSs located on mRNAs and lncRNAs, with 343,185 unique PQSs documented in QUADRatlas [37] and G4Atlas [38] (10nt ≤ length ≤ 60nt). Four sequence identity thresholds were applied: 80%, 85%, 90%, and 95%, corresponding to word size 5, 6, 9, and 10, respectively.

To compare higher dimensional features beyond sequence similarity, 116,846 non-redundant PQSs were prepared using CD-HIT-EST with a sequence identity threshold of 90% to remove duplicated sequences. Applying the same approach to PQSs downloaded from QUADRatlas and G4Atlas resulted in 129,269 unique sequences. In addition, 48,484 mouse piRNA sequences and 16,203 *C. elegans* piRNA sequences (10nt ≤ length ≤ 60nt) were taken from RNAcentral [39] as controls. These four sequence families were sent to RNA-FM [40] and RNAErnie [41] models, where the mean embeddings of each sequence were extracted to represent high-dimensional features. Finally, UMAP was applied to reduce 640-dimensional or 768-dimensional vectors to two dimensions for visualization [42].

### Collection of RNA modification sites

To gain a panoramic view of the RNA modification dataset associated with genes, nine types of modification sites, including m6A, m5C, m5U, m6Am, m7G, m1A, 2’-O-Me, A-to-I editing, and pseudouridine within the exon of the longest transcripts were extracted from the Gene Expression Omnibus (GEO), RMBase database, RMVar database, MiREDiBase, m6A-Atlas database or datasets from the publications [43–46]. We collected data from nine species to compile datasets to construct the database as a proof-of-concept in this pilot study to demonstrate the interplay between RNA modifications and rG4. Modification sites annotated on older genome versions were converted to the latest UCSC or NCBI versions using LiftOver (**Table 1**), including human (UCSC hg38), mouse (UCSC mm39), rat (UCSC rn7), rhesus monkey (UCSC rhemac10), chimpanzee (UCSC panTro5), pig (UCSC susScr11), zebrafish (UCSC danRer11), HIV-1 (NCBI NC_001802.1), and SV40 (NCBI NC_001669.1). Gene annotations of corresponding genomes were downloaded from the Ensembl database release 109. Transcript annotation of mRNA, lncRNA from Ensembl, and microRNA from miRBase v22 were considered [47]. The modification sites of two RNA viruses were based on their genomic RNA or mRNAs.

**Table 1.**
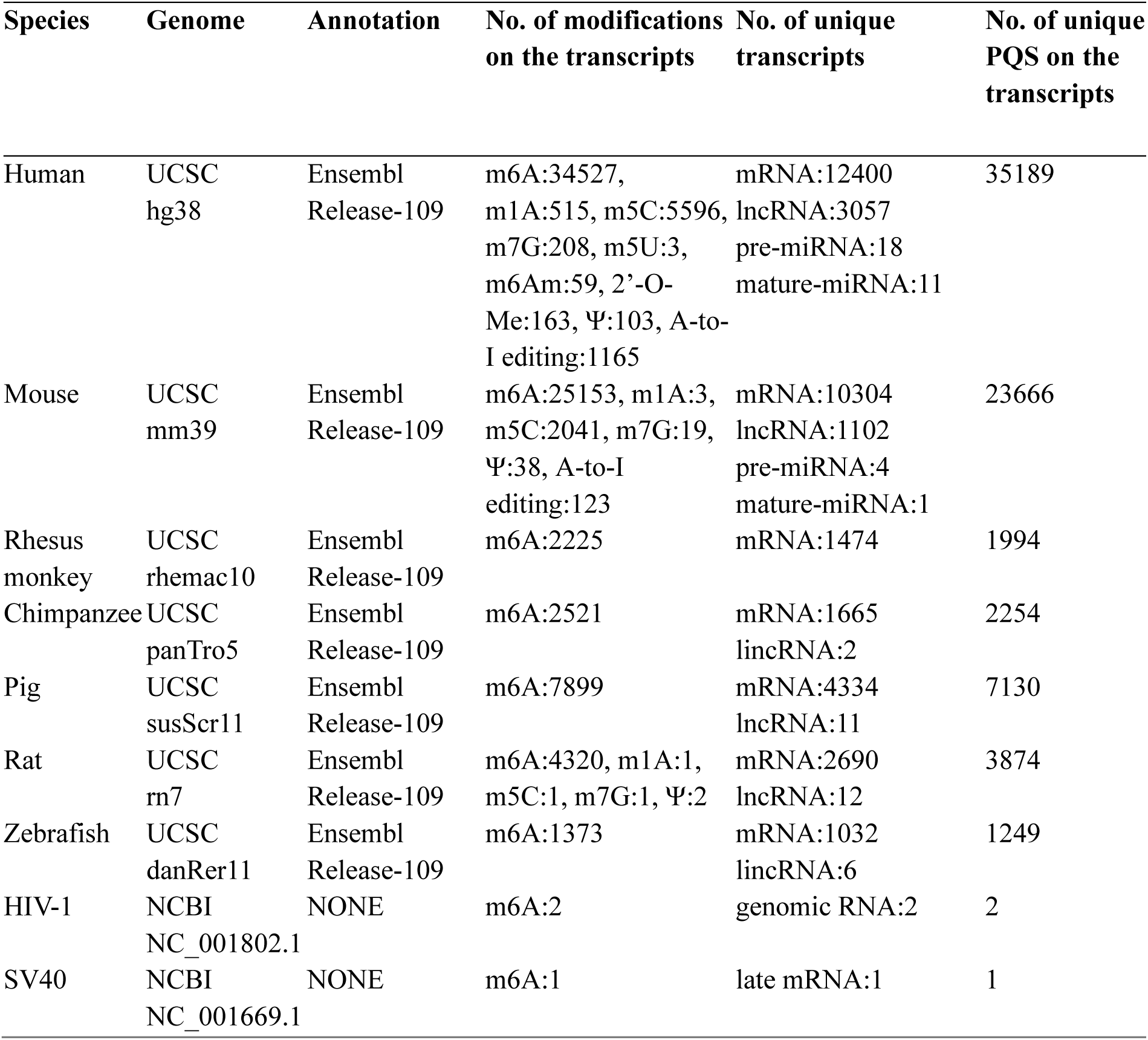
Data source and summary of studied species.

RNA modification sites were categorized into three levels: high, medium, and low according to experimental or predicting methods (**Table S1**). The high level reflects single-nucleotide resolution detection with experimental evidence support, while the low level denotes RNA modifications derived from either databases or predictions. The source of modification and corresponding publication have been incorporated into detailed pages of the MoRNiNG database (**Table S2**).

### Data processing and analysis in the database

RNA modification sites and candidate rG4 loci were processed to obtain base resolution RNA modification sites within rG4 or potential rG4-forming sequences using bedtools [34]. Each modification site was assigned a unique ID following the nomenclature format: species_modification_unique identifier. The location of modifications was further classified into loop, G-tetrad, and bulge to present the modification.

### Database implementation

The database was implemented using SQL Server to hold RNA modifications and PQSs of nine species in independent tables at the backend. The implementation of web page visualization is based on Hyper Text Markup Language (HTML), Cascading Style Sheets (CSS), and JavaScript (JS). Active Server Pages (ASP) scripts were employed to query from the SQL Server backend database through JavaScript by Ajax forms or fetch API. For real-time statistical information, interactive plots were instantly created by ECharts or D3. To display the detailed information of each genomic locus, an embedded JBrowse [48] was inserted into the corresponding sub-pages with a focused view at the selected locus.

## Results

### RNA modification affects the RNA secondary structure that interact with LIN28A

The same RNA segment can form distinct secondary RNA structures depending on its surrounding environment, including the presence of ions and small molecular ligands. While studying the binding sites of RNA binding protein LIN28A, which acts as an m6A anti-reader in mouse embryonic stem cells [17], we discovered that one SL-forming RNA segment identified in its target, *HOXB9*, could also form rG4, as suggested by computational prediction (**Figure 1A, B**). The prediction based on reactivity score, however, only indicated potential SLs **(Figure S1A, B**). Additionally, the profiles captured *in vivo* and *in vitro* did not align well within G-stretches, again deviating from RNAfold prediction. This reflects that RNA structure prediction by RNAfold has limitations to consider factors in solution that could affect RNA folding, such as surrounding ions (**Figure 1C**). In the structure probing datasets, G-stretches are more frequently protected compared to other bases in PQSs, as guanines in G-tetrads showed a statistically lower reactivity score than bases in the loop in several cell lines, including H9, HEK293, HeLa, HepG2, and K562 (**Figure S1C**). Although G tends to be more reactive in structure probing assays and the reactivity score reflects the structure average, the scores of G-tetrads are significantly lower than those of G bases in the non-PQS regions (**Figure S1D**), implying G-tetrads are generally protected in PQSs. To confirm the rG4 formation of this *HOXB9* RNA segment, synthesized RNA oligos were examined. A fully mutated sequence disrupting intramolecular rG4 was designed as a negative control. To demonstrate that the rG4 can form from the sequence that also folds an SL, we further designed two mutated sequences, Mut1_GG2UU_ and Mut2_GG2UU,_ based on the original oligo with a replacement of two bases, GG to UU, at the 5’ end and 3’ end, respectively (**Figure 1A**). Indeed, the fluorescence assay using G4-specific ligands NMM and ThT confirmed the formation of rG4s in the presence of K^+^ (**Figure 1D**). In addition, the rG4 formed *in vitro* exhibited a complex composition according to the electrophoresis and fluorescence assay, as this RNA oligo contains G-tetrads for forming two types of rG4. The comparison of rG4s stained by G4-specific ligand ThT and total RNA dyed by SYBR Gold provided strong evidence that both original and Mut1_GG2UU_ sequences formed rG4 (**Figure 1E, Table S3**). The original oligo likely formed several distinct rG4 structures in solution, given that several types of PQSs were predicted (**Figure S1A**) and the Mut1_GG2UU_ resulted in a much stronger signal in the fluorescence assay (**Figure 1D**). The CD and UV melting results of Mut2_GG2UU_ further confirmed the formation of compound rG4 (**Figure S2**). This is consistent with the observation that compound RNA conformation in solution exhibits lower rG4 absorption than pure rG4 conformation [49] (**Figure S3)**.

**Figure 1.**
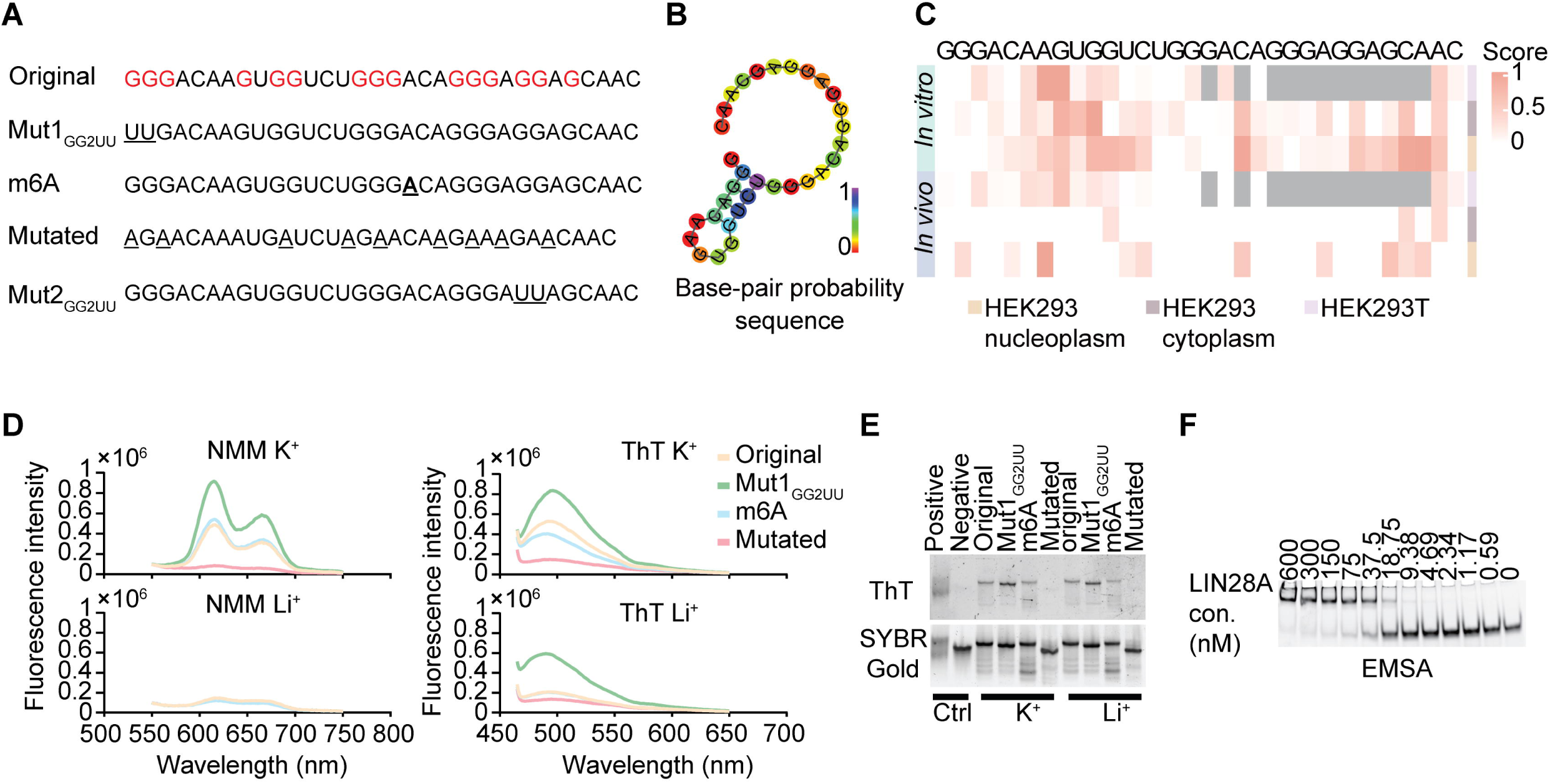
Structural diversity of the *HOXB9* mRNA segment. **A.** Sequences of synthesized oligos of *HOXB9* RNA and its derivatives with mutations and m6A modification. The potential G-tetrads are in red, while UU replacement, m6A, and mutations in the fully mutated sequence are displayed with an underscore. **B.** A predicted canonical SL structure of the original *HOXB9* sequence by RNAfold. The base-pair probability of each base is visualized in color. **C.** The diverse icSHAPE reactivity score of the *HOXB9* RNA segment. The *in vivo* and *in vitro* profiles are captured from HEK293T cells, the cytoplasm and nucleoplasm of HEK293 cells. The score range is from 0 to 1. A higher reactivity score indicates a higher probability of being single-stranded. **D.** The fluorescence assay using G4-specific ligands ThT and NMM under K^+^ and Li^+^ conditions respectively. **E.** rG4 formation of *HOXB9* and its derivatives stained by ThT and SYBR Gold. The conditions to stabilize and destabilize rG4 are considered. The rG4 forming TERRA RNA and G-tetrad mutated TERRA RNA are used as rG4 positive and negative controls, respectively. **F.** EMSA of LIN28A and original *HOXB9* oligo with a gradient concentration of LIN28A in K^+^ condition. The LIN28A concentration ranges from 0 to 600 nM. The concentration of the original *HOXB9* oligo with FAM tag is 10 nM.

As natural m6A modification occurs in this RNA segment, we used a synthesized oligo carrying an m6A modification to investigate rG4 formation. ThT staining demonstrated that the chemically modified adenosine may reduce the rG4 forming potential under the lithium condition that destabilizes rG4 (**Figure 1E**). This reduced stability of rG4 is consistent with the observation in the UV melting experiment, which showed that m6A decreases the thermostability of rG4 (**Figure S4**). The respective rG4 melting temperatures (Tms) of RNA with and without m6A were 55 ℃ and 55.5 ℃ in the forward scan, and 51 ℃ and 53 ℃ in the reverse scan. This suggests that RNA modifications affect rG4 folding.

Next, we examined the binding between rG4 and LIN28A to assess the influence of RNA modification (**Figure 1F, Figure S5**). Despite their binding in the presence of K^+^, we discovered that the RNA carrying m6A showed no significant difference in binding affinity with LIN28A under the rG4 stabilizing condition compared to the RNA without m6A modification. This suggests that m6A potentially affects rG4 under conditions in which rG4 can easily unwind or compete with other conformations. The binding affinity between m6A-carrying RNA and LIN28A indeed declined in the Li^+^ condition (**Figure S5**). The comparison of EMSA under K^+^ and Li^+^ conditions indicates that rG4 enhances RNA binding with LIN28A. LIN28A could also bind to alternative RNA sequences or conformations, potentially influenced by K^+^. This aligns with the observation that m6A can slightly lower the rG4 Tm. This mild change enables conformation to shift toward SL as indicated by the ThT fluorescence assay (**Figure 1D**). The m6A-induced structure change can explain the decreased binding affinity between RNA and LIN28A when rG4 is the preferred binding conformation. Therefore, LIN28A could be captured as an m6A anti-reader in loci where rG4 and SL are nested. These results demonstrate that m6A can affect RNA conformation of rG4-containing sequences and RNA interaction with RBP LIN28A.

### Genome-wide existence of RNAs with multiple-structure forming potential

Rather than recording comprehensive maps of RNA structure dynamics, current RNA structure profiling assays [23, 50–52], including SHAPE-Seq, icSHAPE, DMS-MaPseq, and rG4-seq, primarily provide static snapshots of the structural landscapes in a few experimental settings. It remains largely unknown how distinct secondary structures formed from the same locus transition. We combined computational prediction and public datasets to investigate the potential transition between distinct types of RNA structures. We focused on SL and rG4 for their prevalence and increasingly available experimental datasets. We extracted PQSs from the longest transcripts of mRNA, lncRNA, and miRNA based on consensus PQSs predicted by pqsfinder [27] and QGRS [28], followed by an SL structure analysis using RNAfold in six mammalian and one fish species (**Figure 2A**). A relatively loose threshold was applied to include non-canonical PQSs (see **Methods**). By screening the human and mouse transcriptome, we observed that at least 60% and 50% of PQSs, respectively, can form SL across distinct RNAs. This was predicted by RNAfold based on the minimum free energy (MFE) ensemble (**Figure 2B and 2C**), implying a wide existence of embedded secondary RNA structures. The MFE distribution of canonical PQSs showed a significant difference from imperfect PQSs (**Figure S6**), indicating a relatively flexible nature of imperfect PQSs. Although G-tetrad demands guanine stretches and guanine can pair with uracil in nature [53], the stem of SL is more likely to appear in the loops of PQSs. By analyzing the number and occurrence of PQSs on mRNA and lncRNA in the human transcriptome, the normalized density of PQS counts based on relative positions reveals an uneven distribution of PQSs at both transcript ends. The number of PQSs in each transcript varies significantly, with counts ranging from 1 to 346, suggesting transcripts use rG4 distinctly (**Figure 2B)**. A similar pattern was observed in the mouse transcriptome and other species, except for zebrafish (**Figure 2C** **and Figure S7**). These results imply the widespread existence of secondary structure equilibrium in transcriptomes.

**Figure 2.**
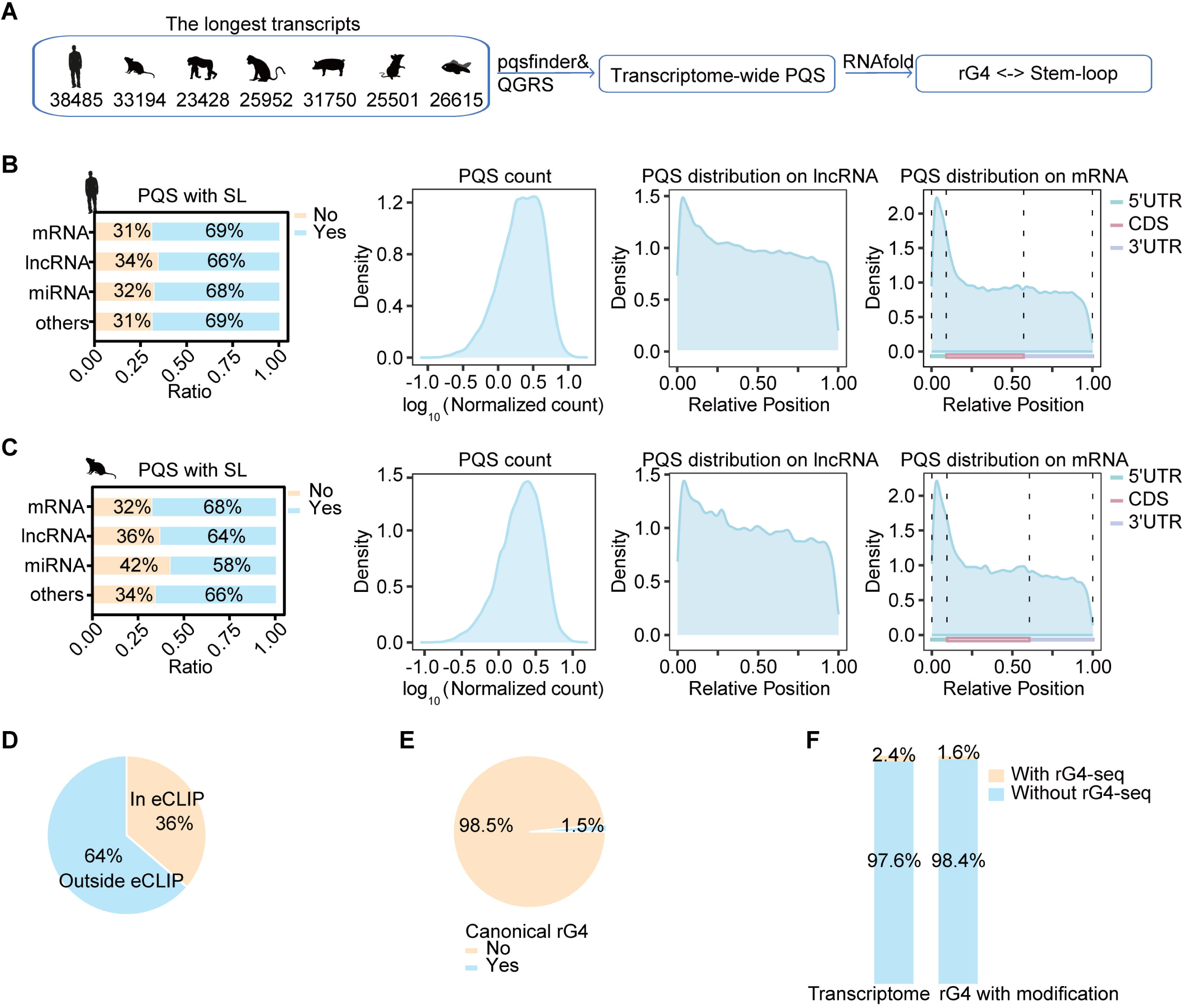
The dual structure-forming nature of RNA segments in transcripts. **A.** Workflow to collect PQSs with SL-forming potential in the transcriptome of vertebrate genomes, including human, chimpanzee, rhesus monkey, pig, mouse, rat, and zebrafish. **B.** The SL-forming potential and distribution of PQSs in the human transcriptome. Left: The detailed proportion of PQSs predicted to form SL in mRNA, lncRNA, miRNA, and the remaining RNAs in the human transcriptome. Middle: The distribution of count per human transcript, normalized to number per kilobases. Right: The relative position of human PQSs in the longest lncRNAs or mRNAs. **C.** The SL forming potential and distribution of PQSs in the mouse transcriptome. Left: The detailed proportion of PQSs predicted to form SL in mRNA, lncRNA, miRNA, and the remaining RNAs in the mouse transcriptome. Middle: The distribution of count per mouse transcript, normalized to number per kilobases; Right: The relative position of mouse PQSs in the longest lncRNAs or mRNAs. **D.** The percentage of PQSs supported by ENCODE eCLIP peaks in human sequences. **E.** The proportion of canonical and non-canonical rG4-forming sequences without SL structure in the human transcriptome. **F.** *In vivo* evidence supported PQSs and modification. Left: The proportion of PQSs supported by rG4-seq in the human transcript. Right: The proportion of PQSs with RNA modifications supported by rG4-seq in the human transcript.

### PQSs and the scarcity of experimental rG4 profiles in the human transcriptome

In the human transcriptome, about 36% of PQSs are supported by at least one eCLIP dataset from ENCODE, consistently demonstrating that RNA binding proteins can interact with rG4 *in vivo* (**Figure 2D).** These PQS-overlapping peaks, which include at least half of the PQSs, were used to identify candidate rG4-binding RBPs. These rG4-binding RBPs were defined as proteins that bind PQS-overlapping peaks, which accounted for at least 15% of the total non-TE peaks in this study. The functional enrichment of rG4 binding RBPs showed a significant overrepresentation in splicing, nucleoprotein granules, and stress granules (**Figure S8**). This result aligns with previous discoveries showing a co-localization between rG4 and ribonucleoproteins [54, 55]. Notably, among the PQSs that RNAfold failed to predict SL structure, we noticed that canonical (G_3_, L_1-7_) and non-canonical rG4 account for 1.5% and 98.5%, respectively (**Figure 2E**). This distribution significantly differed from that of PQSs predicted to form SL. Furthermore, we analyzed the PQSs completely overlapping with rG4-seq peaks or having their last G base located either in the reverse transcription stop (RTS) sites or one nucleotide upstream from rG4-seq and rG4-seq2.0 in the human transcriptome. We found that only about 2.4% of PQSs could be supported by rG4-seq (**Figure 2F**). Additionally, the current rG4-seq dataset solely includes data from HeLa and HEK293T cells, posing a necessity to generate more rG4 profiles to investigate the dynamics of distinct RNA secondary structures. To demonstrate the reliability of PQSs, CD-HIT-EST was employed to cluster PQSs collected in this study with the reference PQSs documented in QUADRatlas and G4Atlas. More than two-thirds (69%) of the reference PQSs were clustered with the collected PQS at a 90% sequence identity threshold; this increased to 87% when the identity was set to 85% (**Figure S9A**). We additionally applied two large language models, RNA-FM [40] and RNAErnie [41], to demonstrate the feature similarity between PQS datasets using two piRNA datasets as outgroups. The two PQS datasets showed good alignment on the projected space after dimensionality reduction, indicating their high similarity (**Figure S9B)**. In contrast, the piRNA sequences from mouse and *C. elegans* formed distinct clusters as expected.

### RNA modifications in rG4 loci

Emerging evidence has shown that the non-canonical RNA secondary structure rG4 is tightly coupled with RNA modifications. Our results further support the idea that RNA bases carrying chemical modifications can alter the base pairing capability of nucleotides in cells to impair the RNA structure equilibrium, as many PQSs can also fold into SL structures. Thus, we collected RNA modification sites in transcripts, including mRNA, lncRNA and miRNA, based on the resources cataloged in GEO, RMBase database, RMVar database, MiREDiBase, m6A-Atlas database, or datasets from the publications, which are found within 15% and 13% of total PQS in human and mouse transcriptomes, respectively. The high-confidence m6A sites within PQSs, supported by experimental evidence at single-base resolution, represent 41% and 32% of the total PQSs that contain m6A sites in human and mouse transcriptomes, corresponding to 14,137 and 7800 sites, respectively. Among human PQSs with modifications, only 1.6% were supported by rG4-seq (**Figure 2F**). To facilitate a systematic investigation of the role of RNA modifications in regulating RNA structure, we compiled the RNA modifications associated with rG4 into an open-access database named **mo**dification of **RN**As **i**n **n**atural r**G**4 (MoRNiNG).

### The MoRNiNG database reports RNA modification loci within rG4 regions

In the MoRNiNG database, we reported RNA modification loci within rG4 validated by experiments or PQSs predicted by several state-of-the-art rG4 predicting tools. We collected nine types of RNA modifications from nine species to demonstrate the preference for modified sites occurring in the transcriptome (**Figure 3A, B;** **Table 1**). The primary function of this database is to host RNA modification loci with tiered reliability for researchers to explore the relationship between RNA modifications and rG4s, and disclose potential biological impacts. To our knowledge, this is the first data collection to build links between canonical and non-canonical RNA structures in natural RNAs. This work also proves that RNA modifications can alter rG4 forming propensity, which may play a role in post-transcriptional regulation. All processed data in MoRNiNG are freely accessible.

**Figure 3.**
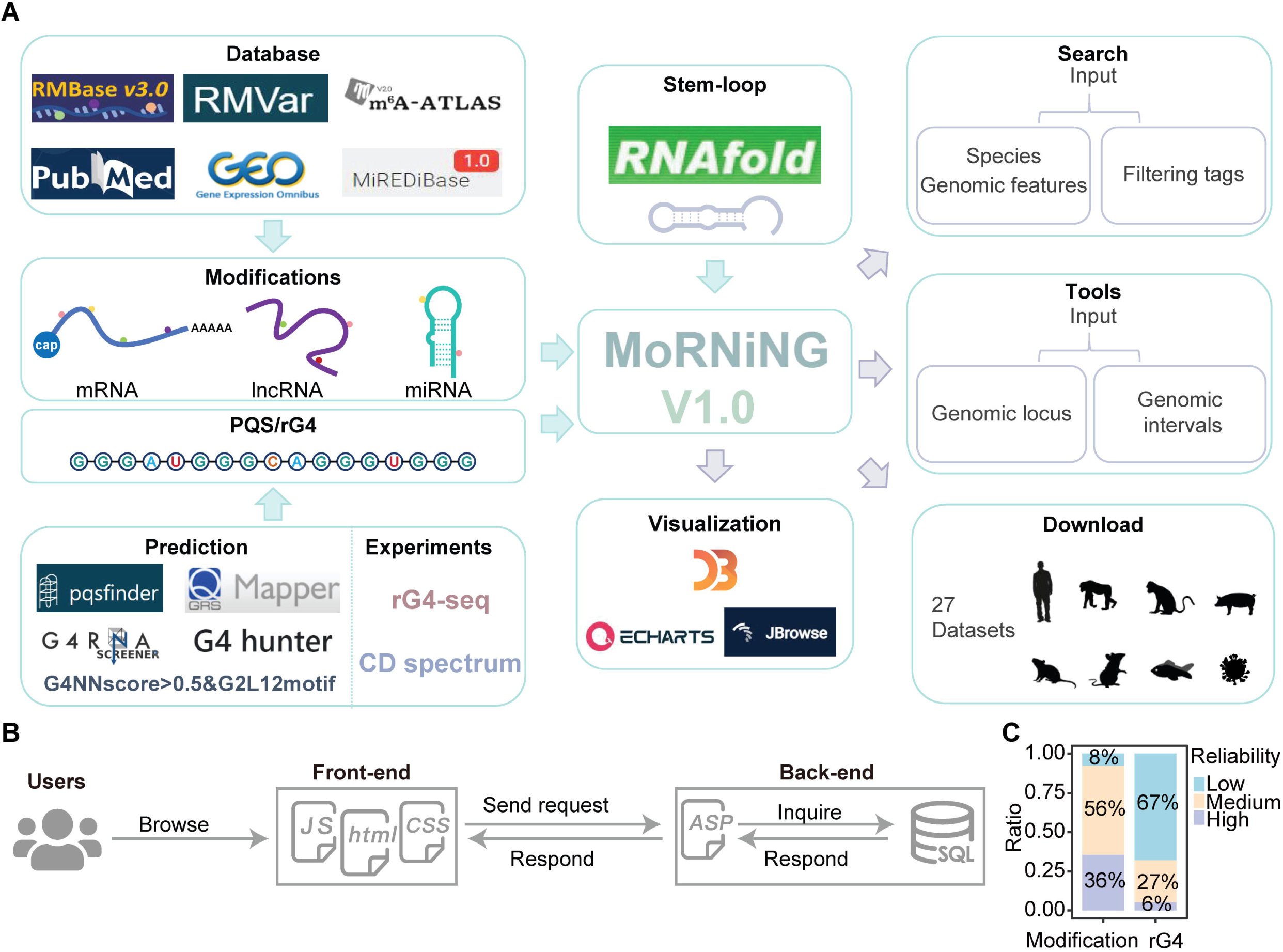
The architecture of the MoRNING database. **A.** A schematic illustration of the data source, approach, and function of the MoRNING database. The source of RNA modification, tools, and experiments for rG4 and PQS, while the major functions, including search, are presented in the center and on the left. The tool and download are summarized on the right. **B.** The architecture and modular implementation of the frontend and backend. The major techniques used to implement the back-end and front-end functions are depicted in groups. **C.** The reliability level of RNA modification sites and rG4/PQSs cataloged in the database.

The MoRNiNG homepage features a real-time tree plot visualizing the constituent species and the number of RNA modification sites cataloged in the database (**Figure 4A**). The positional preference plot of modification sites for the most pronounced modifications in each cataloged species is also visualized. We have implemented three primary pages, including search, download, and tool (**Figure 4B** **and Figure S10**), to help users navigate to distinct functions. A help page was offered to provide instructions, updates, and information on the corresponding data source.

**Figure 4.**
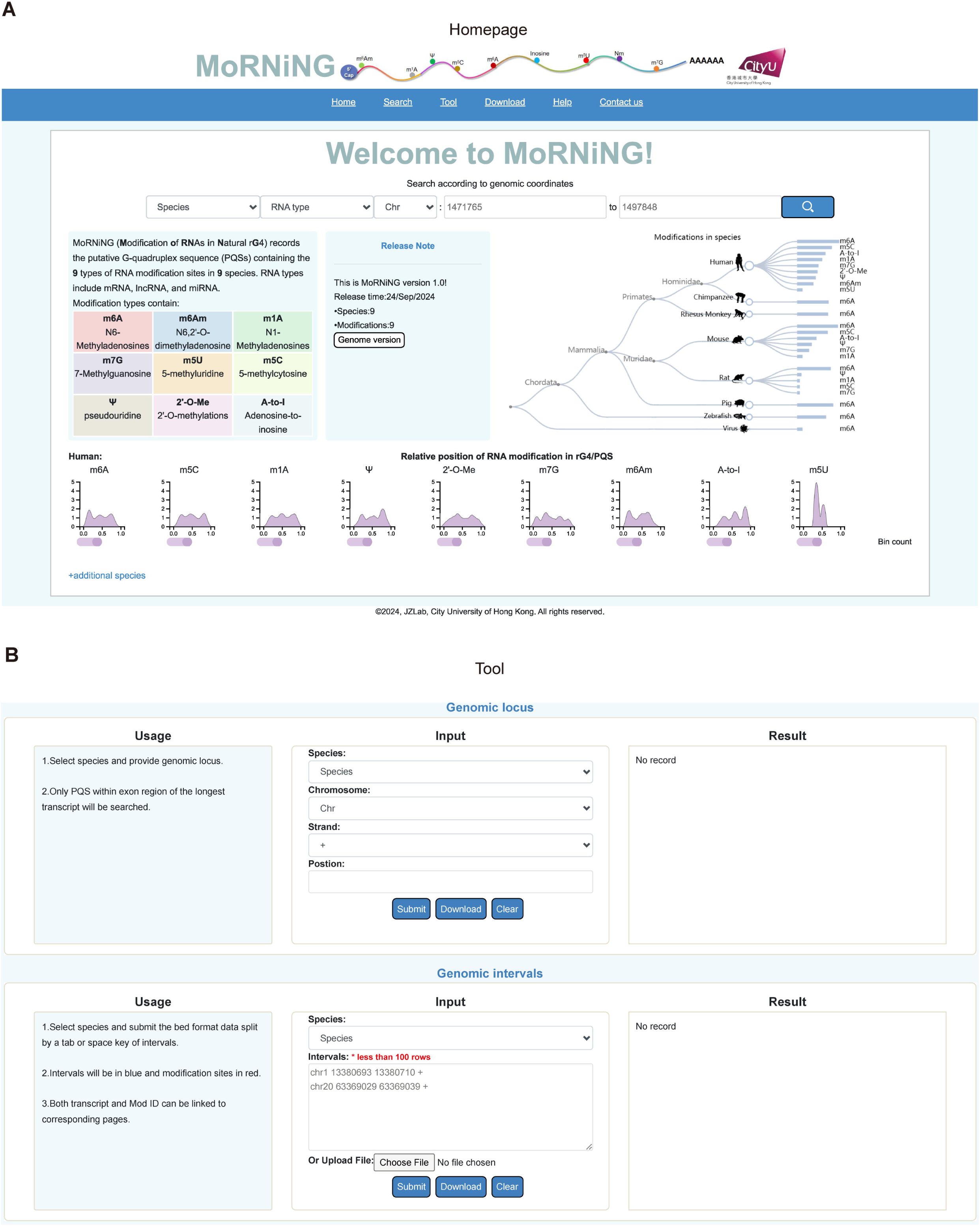
The interface and major functions of the database. **A. and B.** The snapshot of the main page and the tool function.

### Browsing interface

The main page of MoRNiNG database provides a summary exhibiting the total number of RNA modifications in PQSs of the studied species. A search engine is embedded to allow access to the database by specifying the corresponding species and genomic locus via modern web browsers. The search page provides multiple filters under eight categories, including species, chromosome, modification type, RNA type, and four additional options. Users can combine different filtering tags within each species to retrieve desired RNA modifications associated with rG4. The tool page offers two major functions that allow users to upload genomic loci or regions of interest with RNA modifications in a locus-specific or batch mode. Within the download page, formatted table files of RNA modifications for each species can be downloaded for other purposes.

### Tiered reliability and interactive visualization

The RNA modification loci identified by various studies present distinct levels of reliability. Therefore, we classified sites of RNA modifications into three reliability tiers based on the resolution of the corresponding experimental evidence or predictions. Loci in the high-reliability tier are supported by experiments at single-nucleotide resolution. Methods detecting regional RNA modifications are classified as medium, while sites that are fully predicted or from the DARNED [56], RADAR [57, 58] databases are assigned low reliability. PIQS, derived from pqsfinder prediction scores of 10 or higher, is assigned a low-reliability label. The reliability information of sites supported by multiple sources is fully presented. Similarly, we introduced a system with tiered reliability to the rG4/PQS, in which loci with experimental evidence or supported by more than two predictions are labelled as high. Loci supported by one additional predicting tool besides pqsfinder and QGRS are classified as medium. In contrast, loci grouped in the lowest tier are either supported by the consensus sequence predicted by pqsfinder and QGRS or obtained through the setting G4NNscore>0.5&G2L12motif (**Figure 3C**). Each RNA modification site includes the modification type, genomic position, strand, gene symbol, ensemble transcript ID, rG4/PQS coordinate, the relative modification position within rG4/PQS, site type, detection method, and data source (**Figure S11**). An interactive genome browser, Jbrowse, is embedded to visualize rG4 position within the genomic feature, and annotated transcript tracks are available.

### Tools for user-provided genomic locus or intervals

MoRNiNG provides user-friendly web tools for preliminary analysis, including the extraction of RNA modification sites from experimentally validated rG4-forming loci and identifying RNA modification-associated PQS. These two simple tools allow users to query provided loci against existing RNA modifications and rG4s to obtain overlapping predicted rG4s or RNA modification sites within the rG4 and/or PQS in the longest transcript of genes (**Figure S12**). Each search job is executed instantly, and results are visualized upon completion of submitted jobs.

### Data download

All processed RNA modifications within rG4 of distinct RNA species can be downloaded through the browse page after applying selection criteria. Comprehensive modification sites and information stored as zip files can be accessed on the downloaded page for each species. Results generated by querying user-provided genomic loci can also be downloaded.

### Various RNA modifications modulate rG4 dynamics

To comprehensively demonstrate the impact of RNA modifications on secondary structure dynamics, we selected four PQSs from human and mouse transcripts carrying three types of modifications that are relatively abundant *in vivo*. Sequences from these loci include m5C in human *MALAT1* supported by rG4-seq [59], m6A in mouse *Xist* identified by K^+^ dependent RT-stop profiling [10], an A-to-I editing site in human *ABHD2* lacking rG4-seq support, and m5C in mouse *Kmt2d* containing multiple m5Cs supported by ultra-low-input rG4-seq. All rG4-forming sequences were documented in the database. For the *MALAT1* oligo, we found that m5C can potentially promote the formation of rG4 multimer or intermolecular rG4s (**Figure 5A** and **B, Figure S13A**). However, the melting temperature could not be reliably determined for *MALAT1* oligos, probably due to the low rG4 concentration. Compared to the *Xist* oligo without m6A, the m6A-containing oligo exhibited a significantly decreased melting temperature in the forward UV melting scan (**Figure 5C** and **D**). The A-to-I editing significantly enhanced the rG4 formation with considerable thermostability (**Figure 5E** and **F**). We further explored the effect of one m5C versus two m5Cs in the mouse *Kmt2d* and found that all tested ligands could form rG4. However, compared to the oligo without m5C, those oligos containing either one or two m5Cs appeared to enhance rG4 multimer or intermolecular rG4 formation. This effect could be attributed to changes in inter-molecular interactions induced by m5C modifications (**Figure S13B, C, D, E,** and **F**). These findings demonstrate that modifications can influence RNA conformation, further confirming the utility of the database.

**Figure 5.**
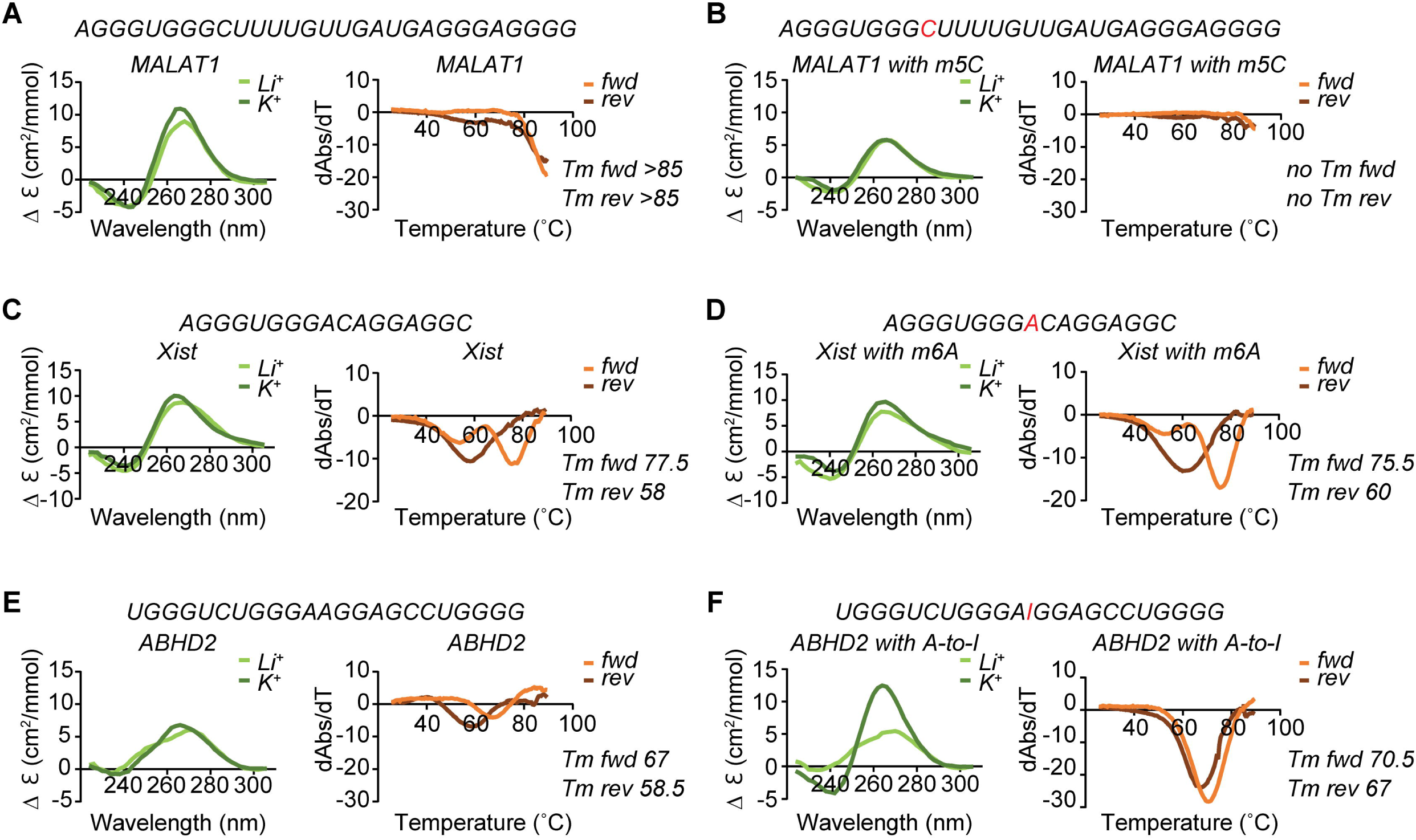
Biophysical characterization of PQSs in *MALAT1*, *Xist* and *ABHD2* and their derivatives with RNA modifications. **A-F**. Left; CD spectrum under K^+^ and Li+ conditions. The CD pattern becomes stronger in the presence of K^+^ rather than Li^+,^ suggesting rG4 formation. The negative peak at 240 nm and the positive peak at 262 nm under K^+^ conditions suggest a parallel topology of rG4. Right: UV melting results. Hypochromic shift observed at 295 nm is a strong sign of rG4 formation. The available Tms for each ligand are indicated. The corresponding sequence of each oligo is shown aside with modifications in red. The forward experiments were conducted from 20 ℃ to 95 ℃, while the reverse experiments were conducted from 95 ℃ to 20 ℃.

## Discussion

The increasing volume of RNA modifications that have been characterized to base resolution poses an opportunity for better understanding the regulatory roles of RNAs, since chemical modification to nucleotides alters the base-pairing or recognition of RBPs. However, relationships between RNA structure and RNA modification remain largely undefined.

Beyond the known SL structure recognized by LIN28A, our experiments demonstrate that the non-canonical secondary structure rG4 can also interact with this protein. The rG4 can facilitate binding with LIN28A, regardless of the presence of potassium. For the existing structure profiling data of LIN28A, poor alignment between the reactivity score and SL can be explained by the formation of rG4 that competes with SL. Indeed, the UV melting experiment shows that m6A can reduce the thermostability of rG4 without drastically impairing the overall binding between RNA and LIN28A. This mechanism coincidentally explains LIN28A binding as an m6A anti-reader, with RNA modification acting as a switch [17]. As the Mut1_GG2UU_ RNA exhibits a lower binding affinity with the protein than the original ligand, we could not rule out the possibility that LIN28A prefers binding a certain form of rG4. Alternatively, a mixture of rG4s could somehow enhance the binding.

The compartmentalized distribution of K^+^ in cells [60] suggests that RNA conformation can dynamically change alongside ions within compartments in response to external cellular stress/stimuli, leading to a shift in structural RNA conformation equilibrium. In addition to the evidence that rG4 and RNA modification are directly coupled [18], RNA modifications are commonly observed in neurons, where ion flux dictates cell excitability. Moreover, several neurodegenerative diseases, including amyotrophic lateral sclerosis, frontotemporal dementia, and Alzheimer’s disease, RNA modifications are directly associated with problematic RBPs and their molecular crowding [15, 16]. The dysregulation of interaction between RNA and RBP suggests a modulation of RNA structure by RNA modifications. The structural changes of RNA also can be regulated by RNA modification, as indicated by several tRNA modifications [13, 61, 62]. The chemical modification of RNA alters the structural stability, which is a potential regulatory layer facilitating RNA structure dynamics. Our findings support a model that RNA modifications, together with the ion environment, can impact the equilibrium of RNA secondary conformation to alter binding affinity with RBPs, thus influencing subsequent regulatory processes. It is also possible that RNA modification can influence intermolecular interactions, as observed in cases with m5C modifications. Although elucidating the detailed mechanism underlying RNA stability shifts remains challenging, the alteration of RNA structure mediated by RNA modification could be used to guide future drug discovery and potentially combined with single-nucleotide polymorphisms to empower personalized medicine treatment strategies.

The large proportion of RNA sequences with dual RNA structure-forming capacity in the genome suggests that the regulation of RNA structures requires greater focus during structure profiling experiments. This also points out another aspect of RNA folding: current RNA folding algorithms require the inclusion of additional parameters to expand the capability to predict and distinguish structural forms derived from the same piece of RNA segment. Our study is limited to *in vitro* assays, providing proof of concept that RNA modification acts as a structure switch by shifting the equilibrium of distinct secondary structures [12]. A similar rule can be potentially extended to the relationship between unstructured RNA segments and structures, as indicated by A-to-I editing. Therefore, a large-scale investigation, *in vivo* profiling, and better RNA folding algorithms would be of great importance to gain a systematic view. However, carefully designed experiments and corresponding simulations demand cross-disciplinary collaborations, which are beyond our current research capacity. Powered by the accumulation of experimental evidence and increased data volume, we established a database to help the RNA research community explore the roles of RNA modification at single-base resolution on their associated rG4s via web browsers. This development opens a door to targeting the regulation of rG4 through RNA modifications. Due to the scope and throughput of extant experimental assays, however, many candidate rG4 loci remain to be validated, and low-confidence loci should be used with caution. Improved rG4-seq protocol and tailored rG4 capturing assays that boost the resolution and throughput would be a valuable resource [26]. Advanced models or algorithms to identify PQS with high accuracy are also on the horizon [40, 41, 63]. Regardless, our study opens a door to targeting the regulation of rG4 through RNA modifications.

In follow-up work, we will focus on several areas: 1) We plan to expand the database capacity by including more RNA modifications from newly sequenced species and from data types produced by other experiments to annotate PQS, such as icSHAPE [20] and HTR-SELEX [6]. 2) We also plan to include or develop a state-of-the-art algorithm to improve the precision of rG4 prediction. Meanwhile, new tools will be introduced to integrate RNA structure data to expand the confidence level of experimental evidence. 3) Through work to automate the data processing pipeline, we will be able to implement a robust data curation procedure, which can quickly include the latest experimental evidence and allow users to submit the published datasets. Overall, the MoRNiNG database will be devoted to providing insights into the interplay between RNA epigenetics and structure dynamics.

## Data availability

MoRNiNG is a database hosting for RNA modifications associated with rG4, which is freely available online. All data can be accessed at https://www.cityu.edu.hk/bms/morning.

## CRediT author statement

**Yicen Zhou:** Data curation, Methodology, Investigation, Visualization, Software, Writing— original draft**. Shanxin Lyu:** Methodology, Investigation, Visualization, Writing – original draft. **Shiau Wei Liew:** Methodology, Investigation, Writing – review & editing. **Xi Mou:** Methodology, Investigation, Writing – review & editing. **Ian Hoffecker:** Investigation, Writing – review & editing. **Jian Yan:** Investigation, Writing – review & editing. **Yu Li**: Investigation, Writing – review & editing. **Chun Kit Kwok:** Conceptualization, Supervision, Investigation, Writing –review & editing. **Jilin Zhang:** Conceptualization, Investigation, Supervision, Investigation, Writing – original draft, review & editing. All authors have read and approved the final manuscript.

## Acknowledgments

We thank Dr. Lisheng Zhang for providing constructive suggestions. The work was supported in part by a start-up grant for new faculty [7200735] and APRT [9610580] to Jilin Zhang by City University of Hong Kong; Research Grants Council of the Hong Kong SAR, China Project [General Research Fund, 11101022] to J.Y; National Natural Science Foundation of China Projects [32471343, 32222089]; Research Grants Council of the Hong Kong SAR, China Projects [CityU 11101525, RFS2425-1S02, CityU 11100123, CityU 11100222, CityU 11100421]; Croucher Foundation Project [9509003]; State Key Laboratory of Marine Environmental Health Seed Collaborative Research Fund [SCRF/0037, SCRF/0040, SCRF/0070]; and City University of Hong Kong projects [9680376, 7030001, 9678302] to C. K. K; and the Hong Kong PhD Fellowship Scheme to S.W.L.

## Supplementary material

Supplementary material is available at *Genomics, Proteomics & Bioinformatics* online (https://doi.org/10.1093/gpbjnl/qzaxxxx)

