## Supplemental Figure Legends for "Modification of RNAs in natural rG4 (MoRNiNG), A Database of RNA Modification Sites Associated with the Dynamics of RNA Secondary Structures"

**Supplementary material**

**Figure S1**	**Computational PQS in the *HOXB9* RNA segment and the icSHAPE reactivity score distribution**

**A.** Predicted PQS within *HOXB9* mRNA by pqsfinder, QGRS, G4Hunter, and G4RNA screener. Potential G-tetrads within the PQS are marked in blue and red, with those suggested by pqsfinder and QGRS underscored. The cGcG, G4H, and G4NN scores in the G4RNA screener are separated by commas. **B.** The predicted canonical SL structure of the original *HOXB9* oligo by RNAfold according to icSHAPE scores in the RASP database. **C.** The icSHAPE score distributions in PQS with or without SL from H9, HEK293, HeLa, HepG2, and K562 cells. G-tetrad-associated bases, and the remaining bases were grouped for each sample. The dot represents the median. The number of PQS in each cell line is shown in the center. The score difference between groups was examined by the Mann-Whitney test. Statistical significance was defined as follows: **P* < 0.05, ***P* < 0.01, ****P* < 0.001, and *****P* < 0.0001. **D.** The icSHAPE score distributions of G bases in PQS and non-PQS from H9 and HeLa cells. The dot represents the median. The number of PQS or non-PQS from each cell line is indicated above the plot for each category. The score difference between groups was examined by the Mann-Whitney test. Statistical significance was defined as follows: **P* < 0.05, ***P* < 0.01, ****P* < 0.001, and *****P* < 0.0001.

**Figure S2**	**Biophysical characterization of the Mut2_GGtoUU_ oligo**

CD spectra and UV melting of the *HOXB9* last G-tetrad mutant sequence. The sequence of the Mut2_GGtoUU_ oligo is shown at the top. The left panel shows the CD spectra of *HOXB9* Mut2_GGtoUU_ oligo in K^+^ and Li^+^ conditions, respectively. The right panel shows the UV-melting of *HOXB9* Mut2_GGtoUU_ oligo from 20 ℃ to 95 ℃ (forward) and 95 ℃ to 20 ℃ (reverse). A hypochromic shift is observed at 295 nm. Both forward and reverse experiments’ Tm value is indicated. fwd: forward. rev: reverse.

**Figure S3**	**The CD spectroscopy of *HOXB9* oligo and its derivatives under K^+^ and Li^+^ conditions**

A stronger signal at 262 nm with a lower signal at 240 nm under K^+^ compared to Li^+^ indicates the formation of rG4 in the original, Mut1_GG2UU_, and m6A oligos. In comparison, the indistinguishable signal at 240 nm and a weaker signal at 262 nm suggest no rG4 is present in the mutated oligo. The negative peak at around 240 nm and the positive peak at around 262 nm under the K^+^ condition suggest a parallel topology of rG4.

**Figure S4**	**The** **UV melting of *HOXB9* oligo and its derivatives under K^+^ and Li^+^ conditions**

The rG4 formation is indicated by a hypochromic shift at 295 nm. The melting temperatures (Tm) derived from the forward experiment in K^+^ buffer are 55.5 °C and 55.0 °C for the original and m6A oligos, respectively. The corresponding Tm values observed in the reverse experiment are 53.0 °C and 51.0 °C. fwd: forward. rev: reverse.

**Figure S5**	***In vitro* interaction of LIN28A protein with HOXB9 oligo and its derivatives**

**A.**	EMSA assay showing the binding of LIN28A protein to the original, Mut1_GG2UU_, m6A, and mutated oligos under Li^+^ condition. **B.** EMSA assay showing the binding of LIN28A protein to the Mut1_GG2UU_, m6A, and mutated oligos under K^+^ condition. The concentration of each oligo is 10 nM.

**Figure S6**	**The MFE distribution of PQSs in human transcriptome**

The database documented PQSs obtained from the human transcriptome are categorized into perfect (G_≥_3, L_1-7_) and imperfect types. The MFE was calculated only for PQSs containing an SL.

**Figure S7**	**Density plots of PQS information in transcripts**

Distribution and number information of PQS on transcripts in the whole transcriptome of **A.** rat, **B.** pig, **C.** rhesus monkey, **D.** chimpanzee, and **E.** zebrafish. Left: the distribution of counts per transcript, normalized to number per kilobases; Right: The relative position of PQS in transcripts.

**Figure S8**	**GO enrichment of rG4-binding RBPs**

Gene Ontology (BP, CC, MF) terms of RBPs in HepG2 and K562 cells for which the ratio of non-TE eCLIP peaks overlapping PQSs to total non-TE eCLIP peaks exceeds 15%. BP: Biological Process. CC: Cellular Component. MF: Molecular Function.

**Figure S9**	**The comparison between collected PQSs and reference PQSs**

**A.** Sequence similarity between PQSs collected in our study and PQSs documented in QUADRatlas and G4Atlas (Reference PQS) using CD-HIT-EST at four identity thresholds. **B.** Shared features of PQSs between datasets. Only non-redundant sequences were subjected to RNA-FM and RNAErnie to extract 640- or 768-dimensional vectors, which were then projected into 2D using UMAP.

**Figure S10**	**Overview of the database interface and main functions**

**A**. Screenshot of the database search page. **B.** Screenshot of the data download interface.

**Figure S11**	**Details of the modification with Mod ID “hsa_m6A_7405”**

The left table contains the information of genomic location, gene and transcript information of “hsa_m6A_7405”, PQS region, PQS length, relative position of “hsa_m6A_7405” on the PQS, site type, and predicted SL structure information. The right table summarizes methods to detect modification and PQS, biological data source, literature source, and genomic coordinates for viewing the PQS in the JBrowse genome browser below.

**Figure S12**	**Identification of an rG4-forming genomic interval on *MALAT1* lncRNA from rG4-seq data using the tool function**

**A.** Querying a genomic interval containing an rG4 on *MALAT1* lncRNA from rG4-seq data. **B.** Output results generated by the query function. **C.** Detailed view of a specific locus with Mod ID “hsa_m5C_4133”.

**Figure S13 Evaluation of rG4 formation of *Kmt2d* and its derivatives**

**(A-C)** CD spectra and UV melting analysis of *Kmt2d* and its derivatives. The sequence of oligo is shown at the top of each panel, with C in red indicating m5C modification. **D**. SYBR Gold and ThT staining assays for *Kmt2d* and its derivatives. **E.** Ligand-enhanced fluorescence assay for *Kmt2d* and its derivatives under K^+^ and Li^+^ conditions, respectively. The emission spectra were recorded from 485 to 650 nm.

**Table S1**	**Approaches of RNA modification detection with credibility and literature sources in nine species**

**Table S2**	**Literature sources of RNA modifications and PQS in nine species**

**Table S3**	**Oligos used for experiments in the study**
