## Supplementary figures and images for "Modification of RNAs in natural rG4 (MoRNiNG), A Database of RNA Modification Sites Associated with the Dynamics of RNA Secondary Structures"

### FigureS1

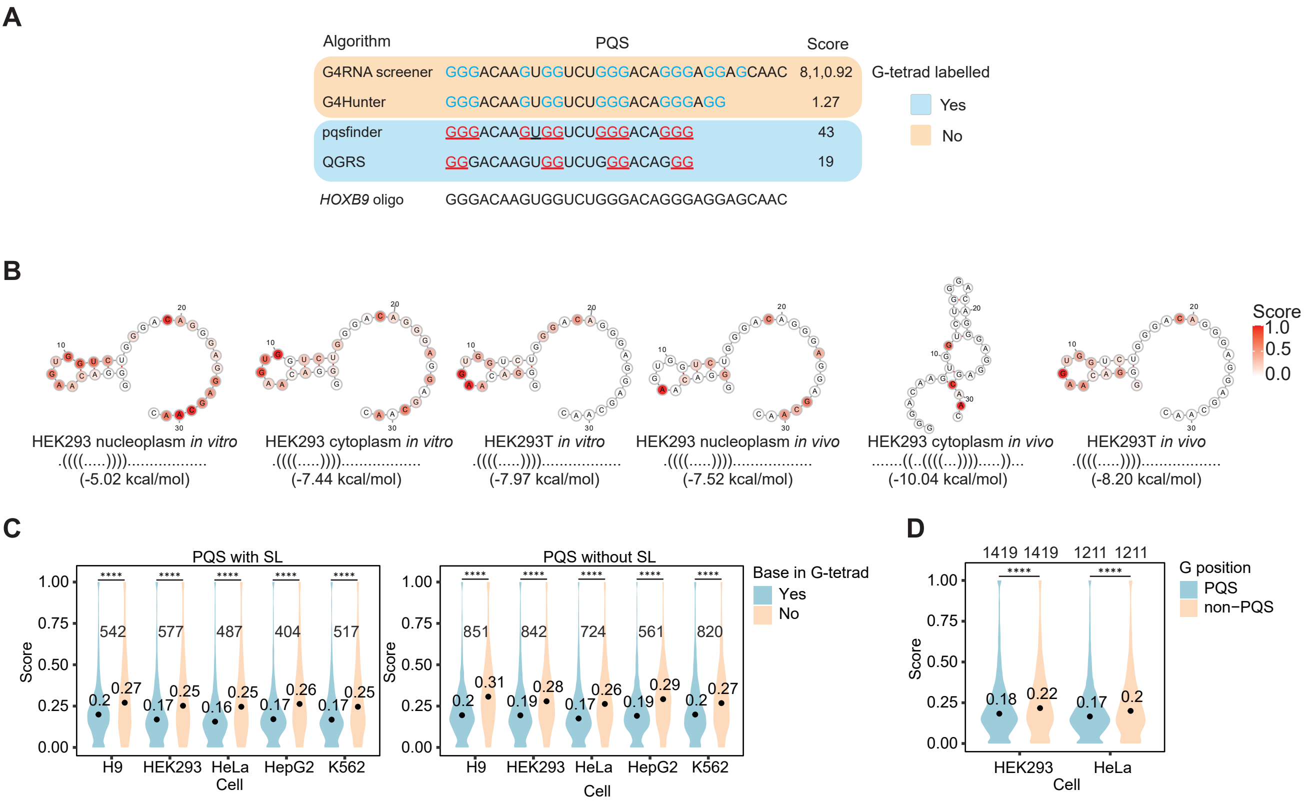

### FigureS2

*GGGACAAGUGGUCUGGGACAGGGAUUAGCAAC*

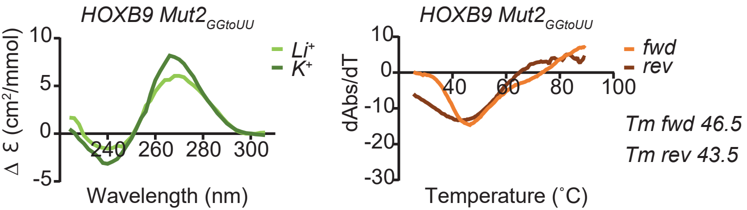

### FigureS3

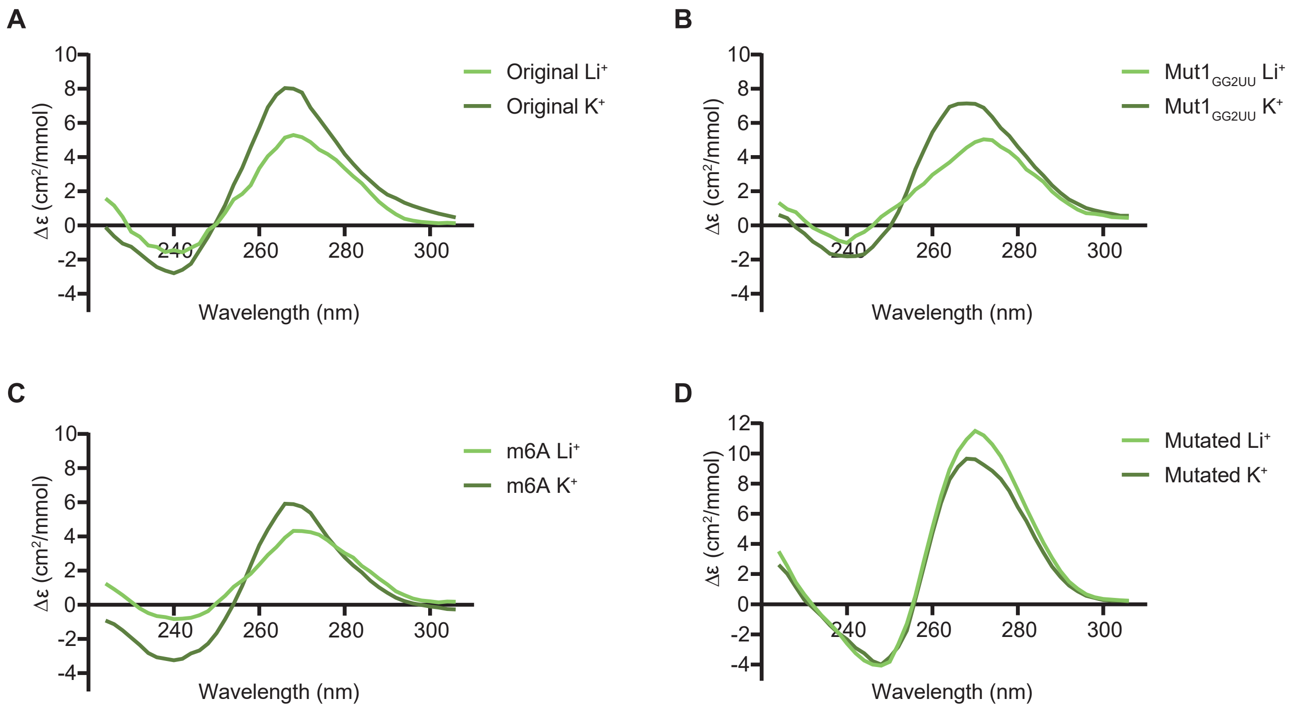

### FigureS4

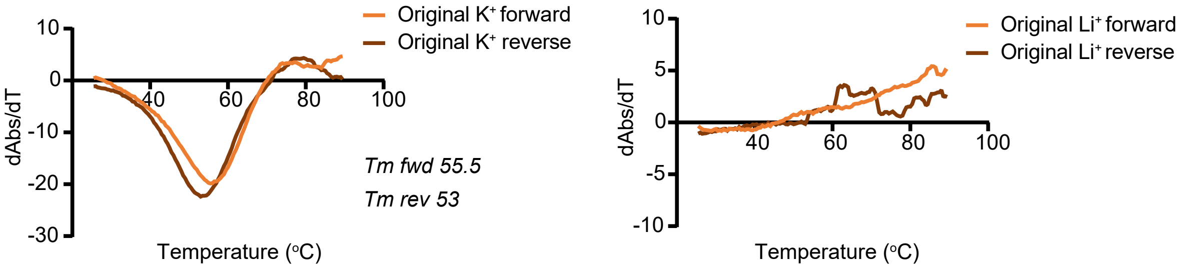

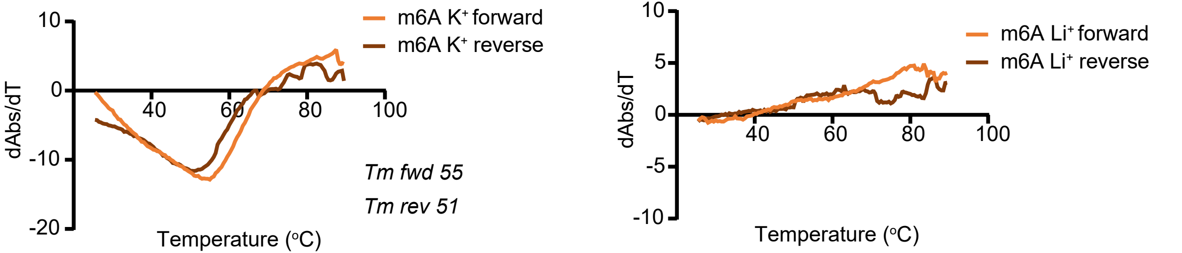

### FigureS5

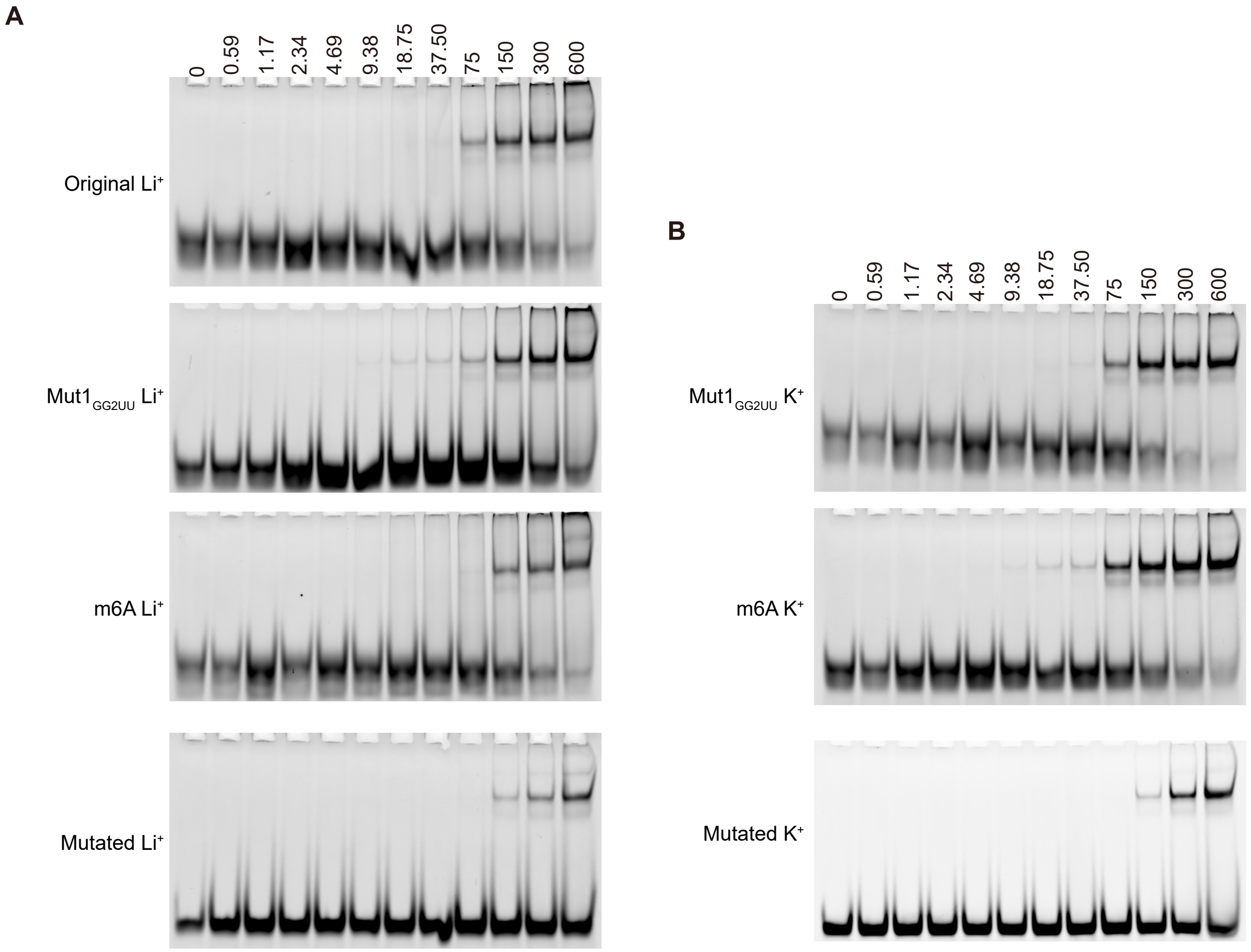

### FigureS6

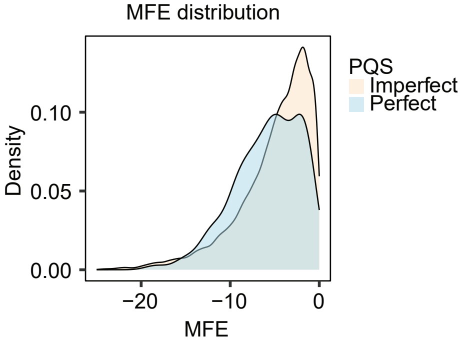

### FigureS7

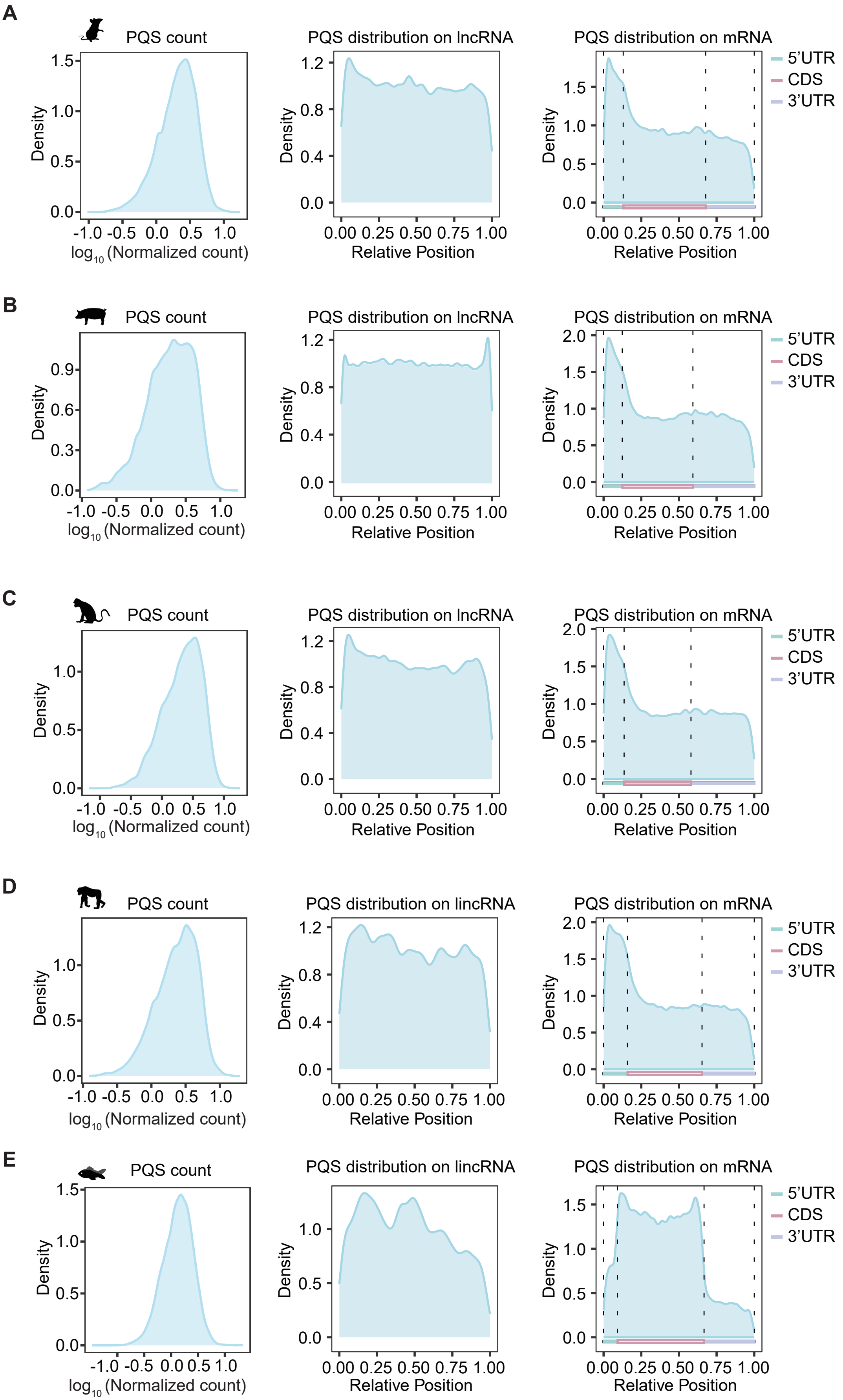

### FigureS8

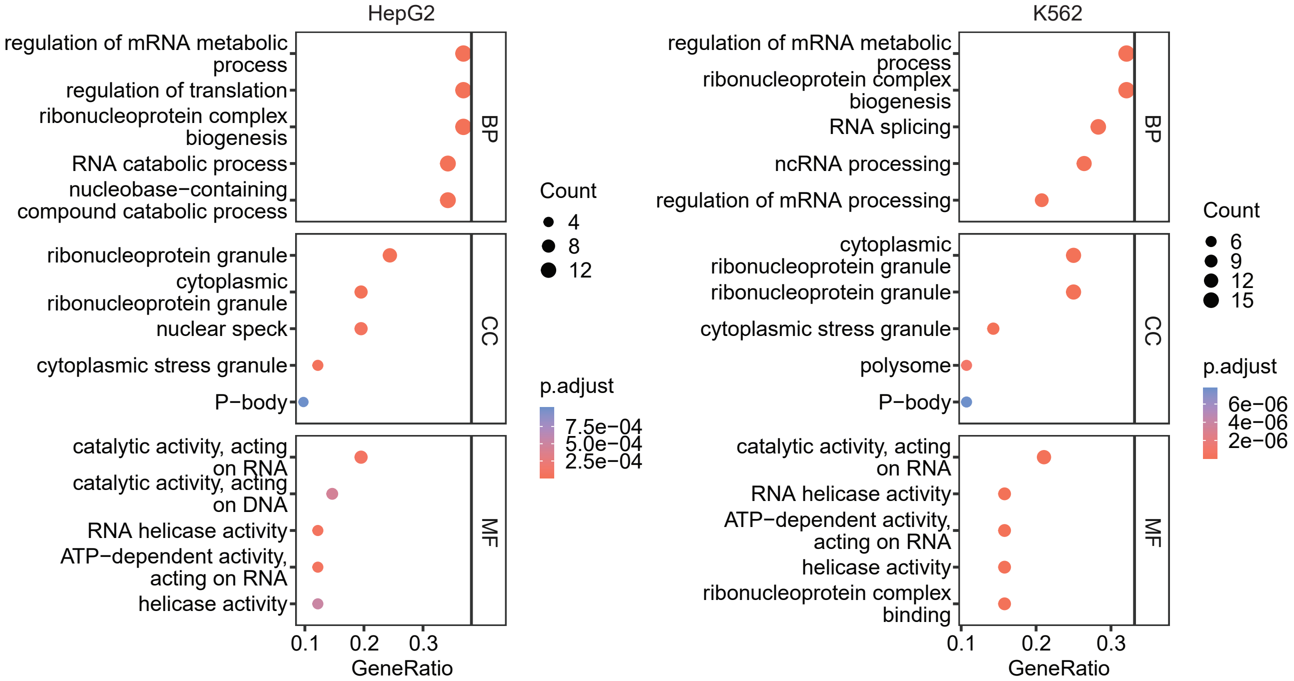

### FigureS9

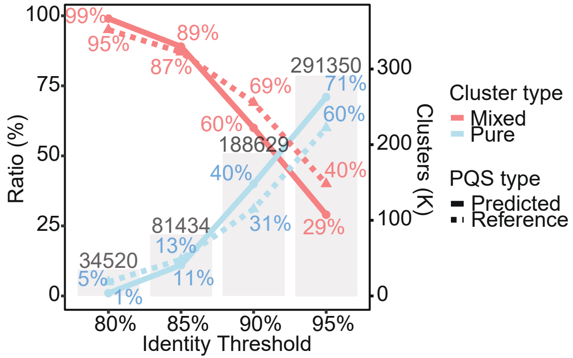

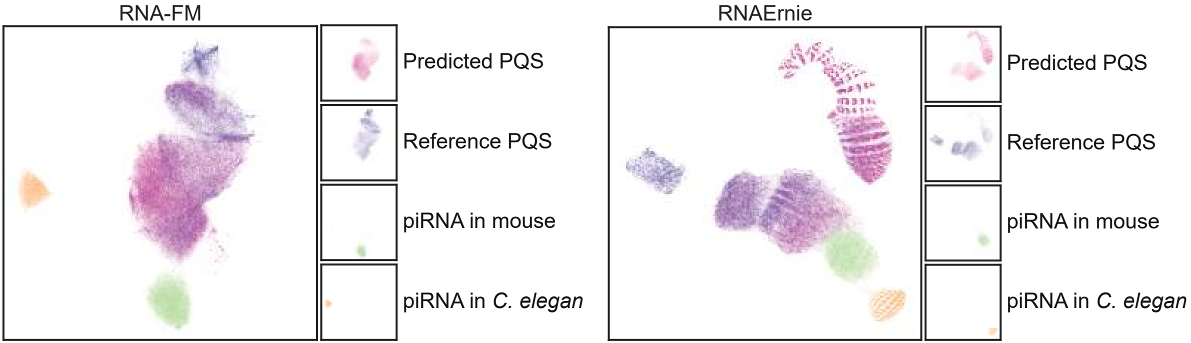

### FigureS12

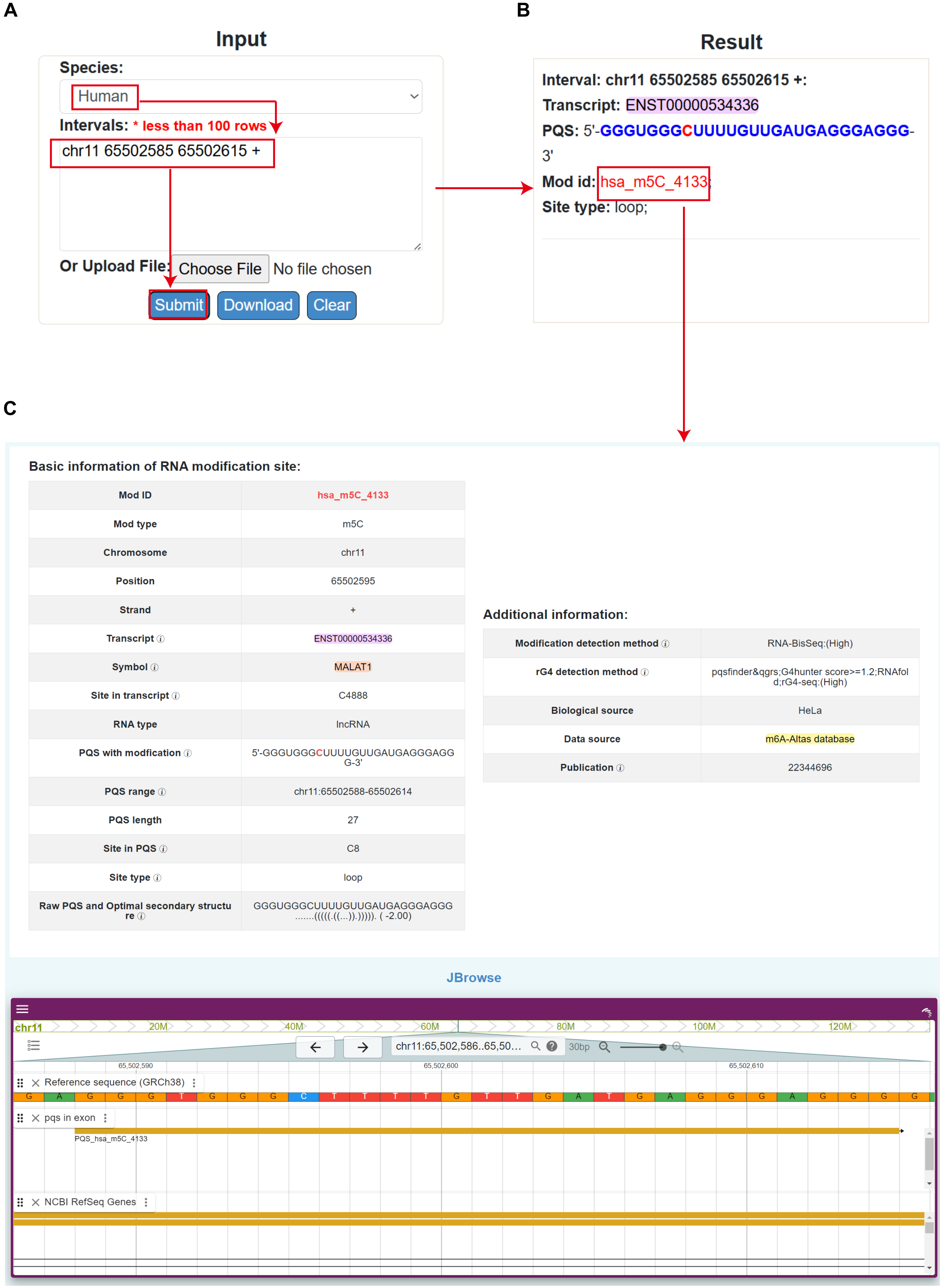

### FigureS13

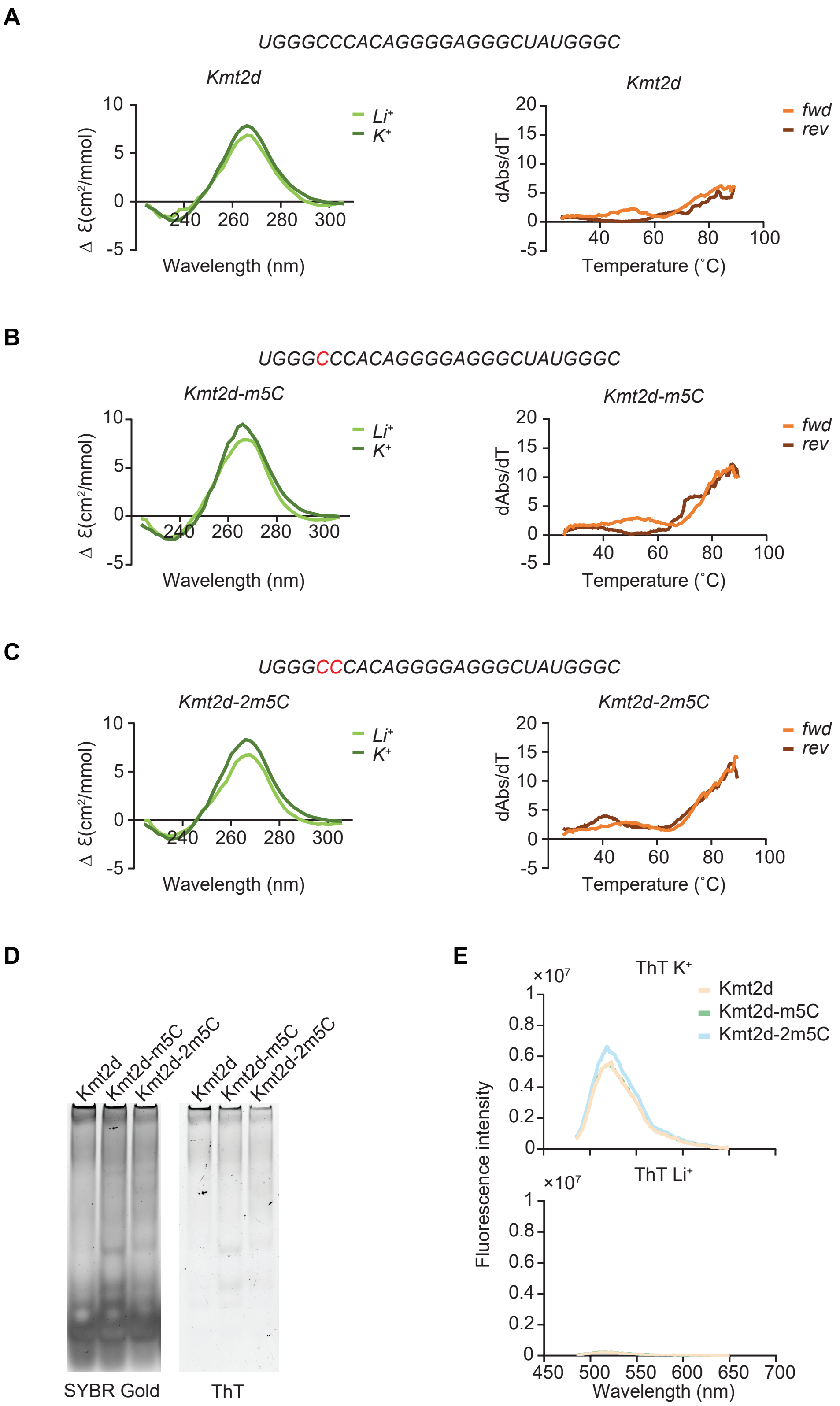
