## Supplementary material for "Modification of RNAs in natural rG4 (MoRNiNG), A Database of RNA Modification Sites Associated with the Dynamics of RNA Secondary Structures": FigureS10

**A**

Search

### Search

Species: Human | Chromosome: chr1 | ModType: m6A | RNA/genome type: lncRNA

Additional Option

csv

Search

| Chr | Position | Strand | ModType | ID | PQS | site in PQS | Transcript | Symbol | RNA/ |
|---|---|---|---|---|---|---|---|---|---|
| chr1 | 1703921 | - | m6A | **hsa_m6A_26600** | 5'-GGCCGUGGACAUGUGGUCAGUGG-3' | A9 | ENST00000598846 | NA | |
| chr1 | 1716390 | - | m6A | **hsa_m6A_26601** | 5'-GGGAAUGGGAAAGACAGAAGAGAAGGGAAAUGGCAAGGG-3' | A14 | ENST00000598846 | NA | |
| chr1 | 151759728 | + | m6A | **hsa_m6A_26748** | 5'-GGCAAGUACUGGAGACAAUGGCUGG-3' | A15 | ENST00000601684 | NA | |
| chr1 | 228274097 | - | m6A | **hsa_m6A_26801** | 5'-GGACAGAGGUGUCUGGAGG-3' | A3 | ENST00000602529 | NA | |
| chr1 | 228274836 | - | m6A | **hsa_m6A_26802** | 5'-GGACCCUCAGGGCAGCAGAGGAUGAGG-3' | A3 | ENST00000602529 | NA | |
| chr1 | 228274532 | - | m6A | **hsa_m6A_26803** | 5'-GGCGGGCGCACAGGACUGG-3' | A15 | ENST00000602529 | NA | |
| chr1 | 228268519 | - | m6A | **hsa_m6A_26804** | 5'-GGCUCUGGGGAUGGACAAGG-3' | A15 | ENST00000602529 | NA | |
| chr1 | 164680135 | + | m6A | **hsa_m6A_26815** | 5'-GGUAGGAUGGACCUACAUGG-3' | A11 | ENST00000602747 | NA | |
| chr1 | 84077916 | - | m6A | **hsa_m6A_26893** | 5'-GGGGGCGGGCGGUGGACCCCCAGCUGGCAGAGCCUGGGGCGGGG-3' | A16 | ENST00000605506 | PRKACB-DT | |
| chr1 | 156642129 | - | m6A | **hsa_m6A_26906** | 5'-GGAGAGGCUGGGGCCGCAGGACAUUCGG-3' | A21 | ENST00000605886 | NA | |

Showing 1 to 10 of 420 rows 10 rows per page ‹ 1 2 3 4 5 ... 42 ›

**B**

Download

### Download

Download by species: Human Zip

Search

| Species | Assembly | DataSet | ModType | Download |
|---|---|---|---|---|
| Homo sapiens | hg38 | human_m6A_info | m6A | Zip |
| Homo sapiens | hg38 | human_m1A_info | m1A | Zip |
| Homo sapiens | hg38 | human_m7G_info | m7G | Zip |
| Homo sapiens | hg38 | human_m5C_info | m5C | Zip |
| Homo sapiens | hg38 | human_m5U_info | m5U | Zip |
| Homo sapiens | hg38 | human_m6Am_info | m6Am | Zip |
| Homo sapiens | hg38 | human_2'-O-Me_info | 2'-O-Me | Zip |
| Homo sapiens | hg38 | human_A-to-I editing_info | A-to-I editing | Zip |
| Homo sapiens | hg38 | human_pseudouridine_info | pseudouridine | Zip |
| Homo sapiens | hg38 | human_PIQS_info | PIQS | Zip |
