## Supplementary material for "Modification of RNAs in natural rG4 (MoRNiNG), A Database of RNA Modification Sites Associated with the Dynamics of RNA Secondary Structures": FigureS11

#### Basic information of RNA modification site:

| | |
|---|---|
| Mod ID | hsa_m6A_7405 |
| Mod type | m6A |
| Chromosome | chr17 |
| Position | 48621735 |
| Strand | - |
| Transcript ⓘ | ENST00000311177 |
| Symbol ⓘ | HOXB9 |
| Site in transcript ⓘ | A2007 |
| RNA type | mRNA |
| PQS with modfication ⓘ | 5'-GGGACAAGUGGUCUGGGACAGGG-3' |
| PQS range ⓘ | chr17:48621730-48621752 |
| PQS length | 23 |
| Site in PQS ⓘ | A18 |
| Site type ⓘ | loop |
| Raw PQS and Optimal secondary structure ⓘ | GGGACAAGUGGUCUGGGACAGGG .((((.....))))......... ( -2.70) |

#### Additional information:

| | |
|---|---|
| Modification detection method ⓘ | GLORI:(High);m6A-seq:(Medium),MAZTER-seq:(High);Prediction by CNN model:(Low);miCLIP2&m6Aboost:(High);CITS_miCLIP:(High);miCLIP:(High) |
| rG4 detection method ⓘ | pqsfinder&qgrs;G4RNA screener:(Medium) |
| Biological source | HeLa,ESC-293T;293T;insulin-treated 293E;HEK293 |
| Data source | RMBase database;34157120;26121403;33756105;m6A-Altas database;RMVar database;36302990 |
| Publication ⓘ | 36302990;26121403;33756105;34157120;31257032;29507755 |

### JBrowse

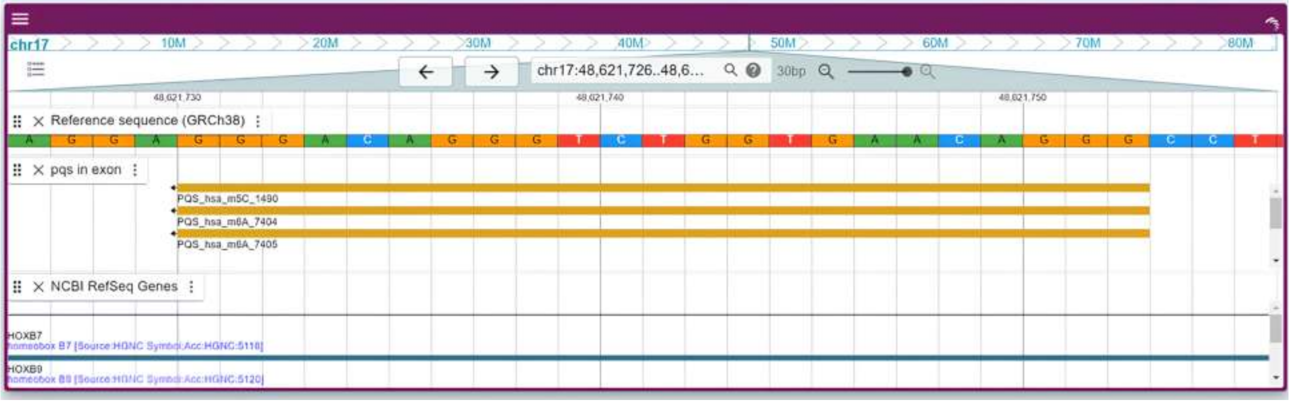
